# Whole-genome and mitogenome based in silico analysis of select *Plasmodium* and identification of a novel drug molecule against the malaria parasite

**DOI:** 10.1101/2022.05.31.494252

**Authors:** Indrani Sarkar, Saurabh Singh Rathore, Gyan Dev Singh, Ram Pratap Singh

## Abstract

Malaria, caused by a parasite known as *Plasmodium*, is one of the leading causes of death, worldwide, especially in the third world countries of Africa and Asia. *Plasmodium* does not only infect humans but also reptiles, birds and other mammals. However, they are the secondary hosts for this parasite. The primary host of *Plasmodium* is female *Anopheles* mosquitoes. Long term researchers have formulated drugs against this deadly pathogen but the current emergence of multi-drug resistance strains of Plasmodium has created a recurring concern. Identification of new drug molecules and understanding their mechanism of action is an urgent need to combat this battle. However, for that, we need to first understand the genomic strategies taken up by Plasmodium to survive the host immune system. With the advancement of high-throughput sequencing techniques, the whole genomes of *Plasmodium* have been sequenced which can help us in advancing our strategies against *Plasmodium*. In this study, we performed a thorough analysis of the genomic features of *Plasmodium* along with its evolution. This revealed a codon biased co-evolution among the parasite and respective hosts. Reverse ecology and protein-protein interactions were studied among *Plasmodium* and *Homo sapiens* revealing a complex biological interaction among them governing the host-parasite interaction as well as drug resistance capability among *Plasmodium*. The molecular docking and simulation studies have found a new drug-target site within mitogenome coding proteins. Those sites were targeted with Cymbopogonol, a phyto-compound derived from *Cymbopogon* (Lemongrass). Along with Cymbopogonol few other *Cymbopogon* derived compounds were also found to be effective as new anti-malarial drug molecules. This is the first report on the effect of *Cymbopogon* derived compounds on *Plasmodium* and is open for a clinical trial.

## Introduction

Malaria, a serious disease which sometimes gets fatal is caused by *Plasmodium*, a parasite which generally infects certain types of mosquitoes (mainly female *Anopheles*) feeding on humans along with several other mammals as well as birds. Patients with malaria generally have a high fever with shaking chills and other flu-like illnesses. According to the World Health Organization (WHO), there were 227 million malaria cases in 85 malaria-endemic countries which increased to 241 million cases in 2020 due to the Covid-19 pandemic and disruption of malaria preventive service distribution due to the pandemic (World Malaria Report 2021: https://www.who.int/teams/global-malaria-programme/reports/world-malaria-report-2021). Although no strong direct relation has been found between the Covid-19 infection and malaria to date, research is still going on to evaluate the effect of one disease upon another (Achan et al. 2022; Wilairatana et al. 2021; Eid, 2021) and the chances for co-infection is always there (Eid, 2021).

Malaria research is now an age-old topic however the advancement in genomics and high throughput sequencing techniques have literally revolutionized this field. Complete genome sequences of several *Plasmodium* species are now available in the public domain database. This has strengthened the speciation strategy of this parasite as well as identified different species-specific hosts also. For example, *Plasmodium falciparum*, *P. vivax, P. ovale*, and *P. malariae* are reported to infect human hosts. Whereas, *P. adleri, P. blacklocki, P. praefalciparum* infect gorillas (Ngoubangoye et al., 2016; Picot and Bienvenu, 2020; Moreno et al. 2013; Galinski et al. 2012, Handunnetti et al. 1987, Schuster 2002). Three species *P. billcollinsi, P. gaboni* and *P. reichenowi* were reported to infect chimpanzees. Macaque monkeys are generally infected by *P. fragile* and *P. cynomolgi* whereas *P. inui* and *P. knowlesi* generally infect rhesus monkeys. *P. berghei, P. yoelii, P. chabaudi* and *P. vinckei* can cause malaria among rodents in general (Ngoubangoye et al., 2016; Picot and Bienvenu, 2020; Moreno et al. 2013; Galinski et al. 2012, Handunnetti et al. 1987, Schuster 2002). Interestingly one species named as *P. gallinaceum* has been reported to infect avian species (Ngoubangoye et al., 2016; Picot and Bienvenu, 2020; Moreno et al. 2013; Galinski et al. 2012, Handunnetti et al. 1987, Schuster 2002). Although there is a host-specific nature persisting among *Plasmodium,* cases of cross contamination are also there establishing malaria as a zoonotic disease (Ramasamy, 2014).

*Plasmodium uses* two different types of hosts in their life cycle – female *Anopheles* mosquitoes as their primary host and humans or other vertebrates as their secondary host. Thus, *Plasmodium* first infects the *Anopheles* mosquitoes and then is transmitted to their secondary host in the form of sporozoites when the mosquitoes bite humans and other vertebrates (Arnot and Gull, 1998). Within vertebrates, there are two separate phases of *Plasmodium* sporozoites - the exoerythrocytic phase which happens in the liver and the erythrocytic phase which happens in the mature erythrocytes or red blood cells (RBC) of the organism. In the exoerythrocytic phase, sporozoites are converted into multinucleated schizonts (pre-erythrocytic stage). Interestingly, Hypnozoites are the quiescent stage in the liver for *P. vivax* and *P. ovale* infection only (Arnot and Gull, 1998). At the end of the exoerythrocytic phase, the schizonts rupture and merozoites are released into the blood and they eventually disrupt the RBC. Within RBC, merozoites mature and form trophozoites and finally multinucleated schizonts. Some merozoites are differentiated into male and female gametocytes. Those gametocytes are ingested by the female *Anopheles* mosquitoes which mature in their mid-gut to form sporozoites and move to their salivary gland ready to be injected into the vertebrates again (Arnot and Gull, 1998). Since there are different stages in the *Plasmodium* life cycle within the secondary host body, different drugs have been developed so far for ceasing the development cycle of this parasite at different stages. For example, Atovaquone-proguanil and primaquine are active against the hepatic-stage schizonts. Primaquine is used against the hypnozoites for *P. vivax* and *P. ovale* infection. Artemisinins, atovaquone-proguanil, doxycycline, mefloquine, and chloroquine are mainly used in the erythrocytic stage as schizonticides (White, 2008). Although several drugs have been developed to date to combat this deadly parasite, we are still at a distance to win over the scenario. One important reason is of course the problem with the health management systems in third world countries. However, another burning factor in the way to triumph in the battle against malaria is the emergence of drug resistance property among *Plasmodium*. There are several reports that this parasite is not only developing resistance against one specific drug but also to a combination of drugs. Moreover, the lack of understanding behind the mechanism of action of combined drug therapies is making the situation worst (Haldar et al. 2018). In this scenario, it is becoming an urgent need to scrutinize the nuclear and mitochondrial genomes of *Plasmodium* and develop new strategies to combat this deadly parasite worldwide.

In this study, we have analyzed 55 *Plasmodium* genomes of different species which are available in the public domain databases. We intended to understand the host-parasite interaction from the genomic and proteomic aspect and tried to identify new drug targets from the mitochondrial genome of *Plasmodium* and utilized the molecular docking and simulation strategies to spot some new plant-derived phyto-compounds against *Plasmodium*. Since *Cymbopogon* sp. (lemongrass) is traditionally used as medicine against fever as well as malaria, we used the *Cymbopogon* derived phyto-compounds to identify some specific component which may prove to be effective against the deadly *Plasmodium* (Chukwuocha et al. 2016).

## Materials and Methods

### Retrieval of whole-genome sequences

The PlasmoDB database (https://plasmodb.org/plasmo/app) was exploited to retrieve the whole genome sequence of 55 *Plasmodium* strains. A detailed list of those select genomes has been given in Table 1. Both the whole nuclear genome sequences of those strains and their respective protein sequences were retrieved. The mitochondrial genome sequences of *Plasmodium* were retrieved from NCBI. We have considered one strain from each species of *Plasmodium.* Moreover, the mitochondrial genomes of some commonly known hosts (both primary and secondary) of *Plasmodium* were also considered. The mitochondrial genomes of primary hosts i.e., *Anopheles stephensi* (NC_028223.1), *A. gambiae* (NC_002084.1), *A. sinensis* (NC_028016.1), *A. funestus* (NC_038158.1), *A. arabiensis* (NC_028212.1), *A. coustani* (NC_050693.1) and *A. merus* (NC_028220.1) were considered. Among secondary host, we have considered *Colomesus psittacus* (NC_015370.1), *Columba livia (NC_013978.1), Falco cherrug* (NC_026715.1), *Falco peregrinus* (NC_000878.1), *Lonchura striata domestica* (CM016794.1), *Passer domestics* (NC_025611.1), *Gorilla gorilla* (NC_001645.1), *Homo sapiens* (NC_012920.1), *Macaca mulatta* (NC_005943.1) and *Pan troglodytes* (NC_001643.1).

**Table 1:** List of select Plasmodium strains considered for this study

| Organism | Genome Version/Assembly ID | Protein coding genes | Orthologs | GO terms |
| --- | --- | --- | --- | --- |
| *Plasmodium adleri* G01 | GCA_900097015.1 | 5325 | 5325 | 3620 |
| *Plasmodium berghei* ANKA | GCA_900002375.2 | 4945 | 4945 | 3666 |
| *Plasmodium billcollinsi* G01 | GCA_900257145.1 | 5317 | 5317 | 3503 |
| *Plasmodium blacklocki* G01 | GCA_900097035.1 | 5100 | 5100 | 3507 |
| *Plasmodium chabaudi* chabaudi | GCA_900095605.1 | 5199 | 5199 | 3715 |
| *Plasmodium coatneyi* Hackeri | GCA_000725905.1 | 5516 | 5516 | 3435 |
| *Plasmodium cynomolgi* strain B | GCA_000321355.1 | 5716 | 5716 | 3274 |
| *Plasmodium falciparum* 3D7 | GCA_000002765.3 | 5318 | 5318 | 4866 |
| *Plasmodium fragile* strain nilgiri | GCA_000956335.1 | 5672 | 5672 | 3454 |
| *Plasmodium gaboni* strain G01 | GCA_900097045.1 | 5134 | 5134 | 3408 |
| *Plasmodium gallinaceum* 8A | GCA_900005855.1 | 5286 | 5286 | 3777 |
| *Plasmodium inui* San Antonio 1 | GCA_000524495.1 | 5832 | 5832 | 3299 |
| *Plasmodium knowlesi* strain A1H1 | GCA_900162085.1 | 4950 | 4950 | 3055 |
| *Plasmodium malariae* UG01 | GCA_900090045.1 | 5942 | 5942 | 4442 |
| *Plasmodium ovale* curtisi GH01 | GCA_900090035.2 | 6671 | 6671 | 3346 |
| *Plasmodium praefalciparum* strain G01 | GCA_900095595.1 | 5812 | 5812 | 4155 |
| *Plasmodium reichenowi* CDC | GCA_000723685.1 | 5638 | 5638 | 4483 |
| *Plasmodium vinckei* brucechwatti DA | GCA_903994205.1 | 5225 | 5225 | 3320 |
| *Plasmodium vivax* P01 | GCA_900093555.2 | 6523 | 6523 | 4504 |
| *Plasmodium yoelii* yoelii 17X | GCA_900002385.2 | 6040 | 6040 | 4591 |
| *Plasmodium cynomolgi* strain M | GCA_900180395.1 | 6068 | 6068 | 2438 |
| *Plasmodium falciparum* 7G8 | GCA_000150435.3 | 5183 | 5183 | 3105 |
| *Plasmodium falciparum* 7G8 2019 | GCA_009761555.1 | 5126 | 5126 | 2528 |
| *Plasmodium falciparum* CD01 | GCA_900617135.1 | 5158 | 5158 | 2455 |
| *Plasmodium falciparum* Dd2 | GCA_000149795.1 | 5209 | 5209 | 2744 |
| *Plasmodium falciparum* GA01 | GCA_900632005.1 | 5224 | 5224 | 2498 |
| *Plasmodium falciparum* GB4 | GCA_900632035.1 | 5335 | 5335 | 2528 |
| *Plasmodium falciparum* GN01 | GCA_900631995.1 | 5298 | 5298 | 2524 |
| *Plasmodium falciparum* HB3 | GCA_000149665.2 | 5186 | 5186 | 2975 |
| *Plasmodium falciparum* IT | GCA_900632055.1 | 5250 | 5250 | 2525 |
| *Plasmodium falciparum* KE01 | GCA_900631975.1 | 5234 | 5234 | 2517 |
| *Plasmodium falciparum* KH01 | GCA_900632025.1 | 5264 | 5264 | 2489 |
| *Plasmodium falciparum* KH02 | GCA_900632015.1 | 5268 | 5268 | 2519 |
| *Plasmodium falciparum* ML01 | GCA_900632085.1 | 5282 | 5282 | 2406 |
| *Plasmodium falciparum* NF135.C10 | GCA_009761425.1 | 5028 | 5028 | 2454 |
| *Plasmodium falciparum* NF166 | GCA_009761515.1 | 5153 | 5153 | 2519 |
| *Plasmodium falciparum* NF54 | GCA_009761475.1 | 5273 | 5273 | 3479 |
| *Plasmodium falciparum* SD01 | GCA_900632095.1 | 5129 | 5129 | 2477 |
| *Plasmodium falciparum* SN01 | GCA_900632075.1 | 5351 | 5351 | 2546 |
| *Plasmodium falciparum* TG01 | GCA_900632065.1 | 5683 | 5683 | 2579 |
| *Plasmodium gaboni* strain SY75 | GCA_001602025.1 | 5198 | 5198 | 3464 |
| *Plasmodium knowlesi* strain H | GCA_000006355.3 | 5328 | 5328 | 3726 |
| *Plasmodium knowlesi* strain Malayan Strain Pk1 A | GCA_002140095.1 | 5259 | 5259 | 3304 |
| *Plasmodium reichenowi* G01 | GCA_900097025.1 | 5644 | 5644 | 3990 |
| *Plasmodium relictum* SGS1-like | GCA_900005765.1 | 5144 | 5144 | 3738 |
| *Plasmodium vinckei* Cameroon EL | GCA_903994265.1 | 5282 | 5282 | 3368 |
| *Plasmodium vinckei* lentum DE | GCA_903994225.1 | 5247 | 5247 | 3342 |
| *Plasmodium vinckei* petteri CR 2020 | GCA_903994235.1 | 5280 | 5280 | 3367 |
| *Plasmodium vinckei* petteri strain CR | GCA_000524515.1 | 5160 | 5160 | 2343 |
| *Plasmodium vinckei* vinckei CY | GCA_900681995.1 | 5051 | 5051 | 3297 |
| *Plasmodium vinckei* vinckei strain vinckei | GCA_000709005.1 | 4954 | 4954 | 2321 |
| *Plasmodium vivax* Sal-1 | GCA_000002415.2 | 5530 | 5530 | 3719 |
| *Plasmodium vivax*-like Pvl01 | - | 5531 | 5275 | 2081 |
| *Plasmodium yoelii* yoelii 17XNL | GCA_000003085.2 | 7724 | 7724 | 2256 |
| *Plasmodium yoelii* yoelii YM | GCA_900002395.1 | 5630 | 5630 | 3928 |

### Characterization of Plasmodium genomes and mitogenomes

A thorough bioinformatic characterization of all the whole genome sequences of *Plasmodium* was performed through CodonW software (http://codonw.sourceforge.net/). We considered total guanine and cytosine (GC) composition, Codon adaptation Index (CAI), Effective number of codons (Enc), and Frequency of Optimal codons (Fop). Genes with the top 10% CAI values were considered to be potentially highly expressed genes (PHX) and genes with the lowest 10% CAI values were potentially lowly expressed genes (PLX). Codons used in the PHX genes were considered to be optimal codons. The index of translational efficiency (ITE) and protein biosynthetic energy cost were calculated by Dambe software (http://dambe.bio.uottawa.ca/DAMBE/dambe.aspx). Statistical correlation analysis (Spearman Rank Correlation) was performed in R software. Total codon and amino acid usage heatmaps were generated in R statistical software. Mitochondrial DNA based phylogenetic analysis was performed through the Baysian algorithm. Finally, the dN/dS based evolutionary analysis was performed on the mtDNA of select *Plasmodium* and their hosts considered for this study.

### Reverse Ecology analysis

Malaria parasites are tremendously adapted to their host’s immune system. They can suppress the human immune system, completely erase the immunological memory for their antigenic determinants and also can alter the structure of the exposed antigens rapidly so that the human immune system gets confused and cannot recognize those determinants. The rapid evolution rate of these parasites is the major reason behind the emerging drug-resistant strains. With the development of bioinformatics tools, it has now become possible to study the population genetics of this malarial parasite to some extent. In this study, we have used reverse ecology, a tool for assessing the interaction between the malarial parasite and the human host. Since a complete whole-genome sequence of *Anopheles* is still not available in the database, we limited our host-parasite interaction study to select *Plasmodium* and the human host. We have used one strain from each species of *Plasmodium* considered in this study.

KEGG Automatic Annotation Server (KAAS) was exploited to retrieve the KEGG ontology of select *Plasmodium* genomes along with *Homo sapiens*. We use RevEcoR (https://github.com/yiluheihei/RevEcoR), an R package to study the competition and complementation index among the aforementioned organisms. This package provides a quantitative measure of interaction in terms of competition index and complementation index. These indices are calculated based on the metabolic needs of the studied organisms. When a group of organisms use the same nutrient sources, the competition (both intra- and inter-species) increases. On contrary, the complementation indices are high for those organisms which are not dependent on the same nutrient source and can co-exhibit without negatively affecting one another. We used R ver 4.1.0. A heatmap based on the competition and complementation index was prepared for better visualization through R software.

### Pathogenic Network analysis

Proteins associated with malaria in both *Plasmodium* and Homo sapiens were taken from the KEGG database (https://www.genome.jp/kegg/). Twenty-four different proteins from *Plasmodium* were solely associated with their pathogenicity (pre-erythrocytic stage and erythrocytic phase). Forty-eight different types of proteins were found to be associated with malaria in humans. The biological network analysis along with the gene ontology assessment of malaria-specific genes from both *Plasmodium* and *H. sapiens* were performed through STRING (https://string-db.org/) and Cytoscape (https://cytoscape.org/).

### Identification of Drug-resistant proteins

ABC transporter family proteins, MSF family proteins and Efflux pump proteins are major protein families associated with the drug resistance of *Plasmodium* (Haldar et al. 2018). We have identified these protein families in all select *Plasmodium* proteomes. We exploited the blastP algorithm for this analysis. Cut-off value was set to be > 98% query coverage and > 98% sequence identity for this analysis. Later we performed a network analysis among those proteins to reveal the actual mechanism of action behind the drug resistance property of *Plasmodium*.

### Selection of mitochondrial proteins as drug targets

The mitochondrial genome is the major source of power supply in organisms. Among *Plasmodium* this organellar genome also plays a crucial role. Previous studies have found that there are some functional and molecular differences between *Plasmodium* and their host mitochondria (Lunev et al. 2016). It has been assessed that; oxidative phosphorylation is not the major function of *Plasmodium* mitochondria in their erythrocytic stage of the life cycle (Rodrigues et al 2010; Ke et al. 2016). During this phase, they rely on glycolysis for energy. The fact that glucose uptake increases by 75-100-fold in Plasmodium-infected RBCs in comparison to non-infected RBCs further validated the importance of glycolysis in *Plasmodium* life cycle (Vaidya and Mather,2009; Bryant et al. 1964). This excess glucose uptake from the blood within the cells results in hypoglycemia as well as increased lactate production. Both of them contribute to lactic acidosis which is the major cause of malaria mortality (Roth et al 1988; Planche and Krishna, 2006). Thus, the main function of mitochondria in *Plasmodium* erythrocytic stage is not oxidative phosphorylation but to maintain the inner mitochondrial membrane potential.

Malarone which is the combination of Atovaquone (mitochondrial bc1 complex inhibitor) and Proguanil (dihydrofolate reductase (DHFR) inhibitor) has been proved to be effective in arresting the growth of the parasite and collapsing its inner mitochondrial membrane potential (Arnot and Gull,1998). Another mitochondrial target which is gaining the attention of the whole scientific world as an anti-malarial drug target is dihydroorotate dehydrogenase (DHODH) (Fleck et al. 1996; Srivastava et al. 1997). In the RSCB-PDB database, we found the crystal structures of dihydrofolate reductase (*P. falciparum*) and dihydroorotate dehydrogenase (*P. vivax*) with PDB ids-2BL9 and 6VTY respectively. Unfortunately, we could not find any crystal structure of *Plasmodium ’s* mitochondrial bc1 complex. We tried to build the protein 3D model of the *Plasmodium’s* bc1 complex however, that approach failed due to the non-availability of proper template proteins. Another interesting fact to be mentioned here is, that the crystal structure with PDB id-2BL9 has pyrimethamine bound to DHFR of *P. vivax*. Pyrimethamine is a well-known anti-malaria drug. However, its usage is now hampered by the emergence of resistance against it among *Plasmodium*.

Since we are more concerned about the drug resistance of this parasite we intended to compare the binding attributes of pyrimethamine and our select ligand compounds (Kongsaeree et al. 2005). The 6VTY was a structure of *P. falciparum* DHODH with a pyrrole-based inhibitor. Thus, it will also be interesting to compare the efficacy of our select compounds against the proven DHODH inhibitor (Kokkonda et al. 2020).

### Preparation of protein structures for molecular docking analysis

Two PDB structures with PDB IDs-2BL9 and 6VTY were downloaded from the PDB database (https://www.rcsb.org/). Those compounds were prepared for the molecular docking analysis after the deletion of all water molecules and the addition of polar Hydrogen molecules. Finally, the Histides were neutralized and Kollman charges were added to the protein structures. All of these aforementioned steps were performed in Autodock MGLtools.

### <u>Cymbopogon</u> derived phytocompounds

*Cymbopogon* sp. (also known as Lemongrass or citronella grass or fever grass) is well-known for its mosquito repellent nature. Recently, it has been proved that lemongrass oil can interfere with the olfactory system of mosquitoes (Maguire et al., 2022). However, the effect of *Cymbopogon* derived phytocompounds against the *Plasmodium* remains largely unknown. Hence, we exploited Indian Medicinal Plants, Phytochemistry and Therapeutics (IMPPAT), a curated database to search all the active compounds present in *Cymbopogon.* A total of thirty-four compounds were found from IMPPAT (Table 2). The structures of those compounds were downloaded in ‘sdf’ format from the NCBI-Pubchem database and were later converted into ‘pdbqt’ format through Open Babel. Those pdbqt formats were used for molecular docking analysis. The draggability of those compounds was assessed by Lipinski’s rules, PAINS and Veber.

**Table 2:** Phytocompounds from Lemon grass and their docking scores with 2BL9 and 6VTY

| Phytocompounds | 6VTY_docking score(kcal/mol) | 2BL9_docking score (kcal/mol) |
| --- | --- | --- |
| Cymbopogonol | -15.1* | -15.7* |
| Triterpenoid | -14.8 | -13.3 |
| Cymbopogone | -13.3 | -15.6 |
| Phytosterols | -10.7 | -13.2 |
| BETA-CARYOPHYLLENE | -10.3 | -11.8 |
| luteolin | -10.0 | -10.0 |
| Sesquiterpene II alcohol diglucoside<br>(Compound 1) | -10.0 | -10.7 |
| Homoorientin | -9.9 | -10.3 |
| p-Camphorene | -9.7 | -10.7 |
| 1-Hexacosanol | -7.6 | -6.3 |
| glutathione | -7.6 | -6.4 |
| (-)-Menthone | -7.2 | -5.8 |
| 3-Carene | -7.2 | -8.7 |
| Eucalyptol | -7.1 | -6.5 |
| dl-Menthol | -7.0 | -7.8 |
| Geranylacetone | -7.0 | -7.9 |
| alpha-TERPINEOL | -6.8 | -7.5 |
| GERANYL ACETATE | -6.8 | -6.3 |
| Dipentene | -6.7 | -8.5 |
| Citral | -6.5 | -7.2 |
| nerol | -6.4 | -7.3 |
| D-Citronellol | -6.2 | -7.0 |
| (-)-Linalool | -6.1 | -5.5 |
| .beta.-Elemol | -6.1 | -7.2 |
| 2-Undecanone | -6.0 | -5.0 |
| GERANIOL | -5.9 | -7.0 |
| CITRONELLAL | -5.8 | -7.0 |
| LINALYL ACETATE | -5.8 | -6.1 |
| MYRCENE | -5.8 | -5.4 |
| 6-METHYL-5-HEPTEN-2-ONE | -5.5 | -6.3 |
| Decanoic acid | -5.4 | -6.0 |
| (E)-6-Methylhept-3-en-1-ol | -5.3 | -5.9 |
| octanoic acid | -5.3 | -5.5 |
| acetone | -3.2 | -2.8 |

### Molecular docking and simulation

Although both 2BL9 and 6VTY were attached to their respective inhibitors and thus their active sites were already identified, we preferred to perform blind docking and then site-specific docking. The idea was to reveal whether the *Cymbopogon* derived phytocompounds bind at the already known active site or a new site. Through this approach, we would be able to search for new drug targeting sites in our select protein structures. After blind docking, we constructed a grid specific to the best binding site and redocked our phytocompounds. Later, the interacting amino acids of *Cymbopogon* derived phytocompounds were compared with already known active sites of both proteins. Molecular dynamics simulation was performed with the best protein-ligand complex for both 2BL9 and 6VTY.

### Molecular dynamics simulation

MD simulation was performed in GROMACS. At first, a cubic box was set up in such a way that the receptor-protein complex will remain away from each side of the box by an equal distance of 1.0nm. Solvent molecule ‘spc216’ was assigned. After preparing the cubic water box, the ionic charge neutralization was done with Na^+^ and Cl^-^ ions. The pressure was set at 1bar and temperature was set at 300K. The simulation was run for 10ns. The RMSD and RMSF fluctuations were compared between the apo-protein (protein without any ligand) and the receptor-ligand complex. MM-GBSA (molecular mechanics generalized Born surface area) was installed within GROMACS and the free energy calculation was performed with MM-GBSA.

## Results

### Characterization of <u>Plasmodium</u> genomes

The *Plasmodium* genomes were highly AT-rich. The potentially highly expressed genes as well as the potentially lowly expressed genes were also AT-rich. The overall AT% varied from 75-77% among the select *Plasmodium*. The third position of the codons was also dominated mainly by A3 (62-68%) followed by T3 (56-58%) rather than G3 and C3. The same trends persisted for both PHX and PLX genes. Moreover, PHX genes were more AT biased (82-85%) rather than PLX (69-72%). Since *Plasmodium* happens to be a highly AT-rich genome, we compared correlated AT3 and ENc. They were significantly negatively correlated (r=-0.65 to -0.69, p<0.001). The CAI and AT were found to be positively correlated (r=0.79 to 0.82, p<0.001). A significant positive correlation was found between Fop, CAI and AT (r=0.85 to 0.89, p<0.001) among *Plasmodium*.

Regarding the codon usage pattern of *Plasmodium* AAU, AAA, GAA, GAU, UUA, AUA, UAU, AUU, UUU, CAA, ACA, AGU, GUU, GGA, UCA, GUA, AAC, AUG, AAG, GGU, CCA, CAU, AGA, GCA, UGU, UCU, ACU, GCU, CCU codons were found to be the most used codons. Among them, AAU, AAA, GAA, GAU, UUA, AUA, UAU, AUU, UUU, CAA were the optimal codons (Fig1). Most interestingly, the AT richness property was also found in the mitochondrial genomes of *Plasmodium*. This indicated an AT dominated compositional constrain among *Plasmodium*. This codon usage pattern was compared with both *Anopheles* and humans. Twenty-eight codons (AAU, AAA, GAA, GAU, UAU, AUU, UUU, CAA, ACA, AGU, GUU, GGA, UCA, AAC, AUG, AAG, GGU, CCA, CAU, AGA, GCA, UGU, UCU, ACU, GCU, CCU, GAG, UUG) were found to be used favourably both in *Plasmodium* and human. Twenty-three codons (AAU, AAA, GAA, GAU, AUU, UUU, CAA, AGU, GUU, GGA, GUA, AAC, AUG, AAG, GGU, CCA, CAU, GCA, UGU, ACU, GCU, GAG, UUG) were preferred between *Plasmodium* and *Anopheles*. A total of twenty-two (AAU, AAA, GAA, GAU, AUU, UUU, CAA, AGU, GUU, GGA, AAC, AUG, AAG, GGU, CCA, CAU, GCA, UGU, ACU, GCU, GAG, UUG) codons were preferred among *Plasmodium*, human and *Anopheles* (Fig 1).

**Fig 1.**
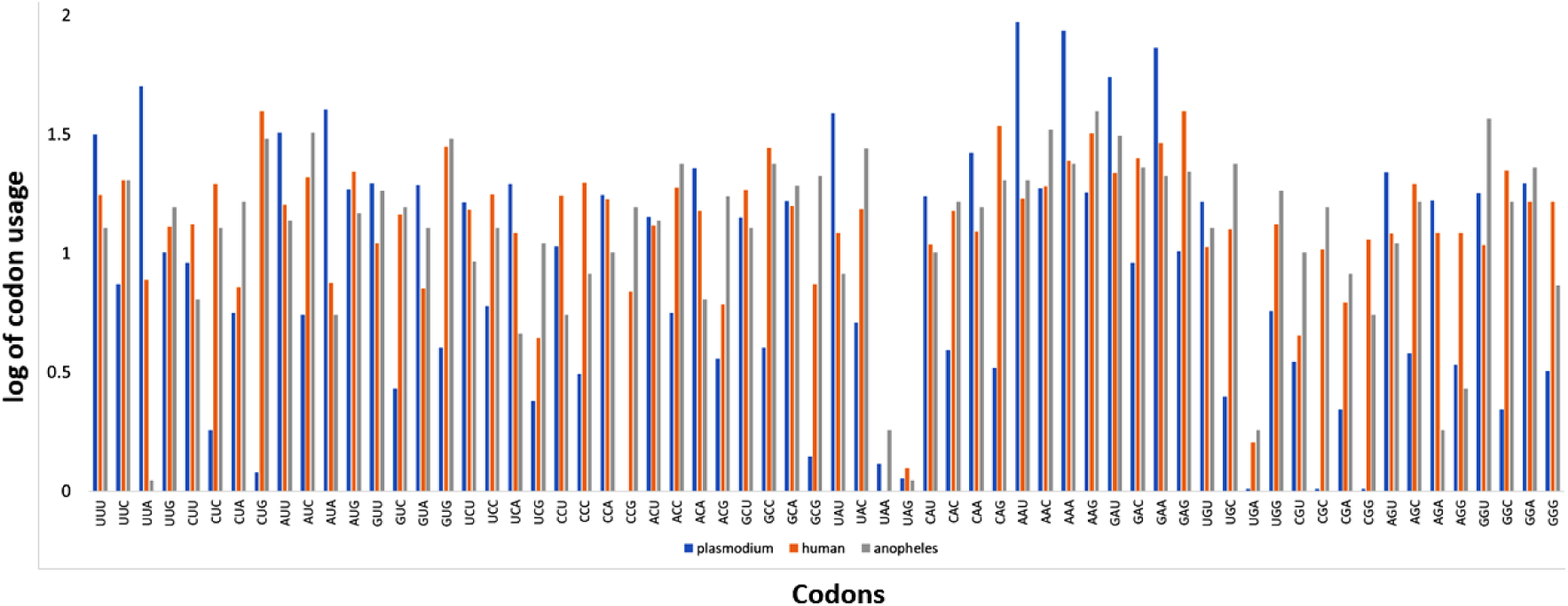
Codon usage comparison among select *Plasmodium*, *Anopheles* and Human revealing a similar pattern of codon bias nature indicating towards codon co-evolution among hosts and parasites.

Regarding the amino acid usage and their synonymous codon usage pattern of *Plasmodium* - Asn (AAU), Lys (AAA), Ile (AUA), Leu (UUA), Glu (GAA), Ser (UCA), Asp (GAU), Tyr (UAU) were found to be exploited more than other standard amino acids (Fig 2). Except, Tyr all other most preferred amino acids are energy economic. Moreover, the frequently used synonymous codons coding those aforementioned amino acids were also AT-rich and mostly dominated by either A or T at the third position.

**Fig 2.**
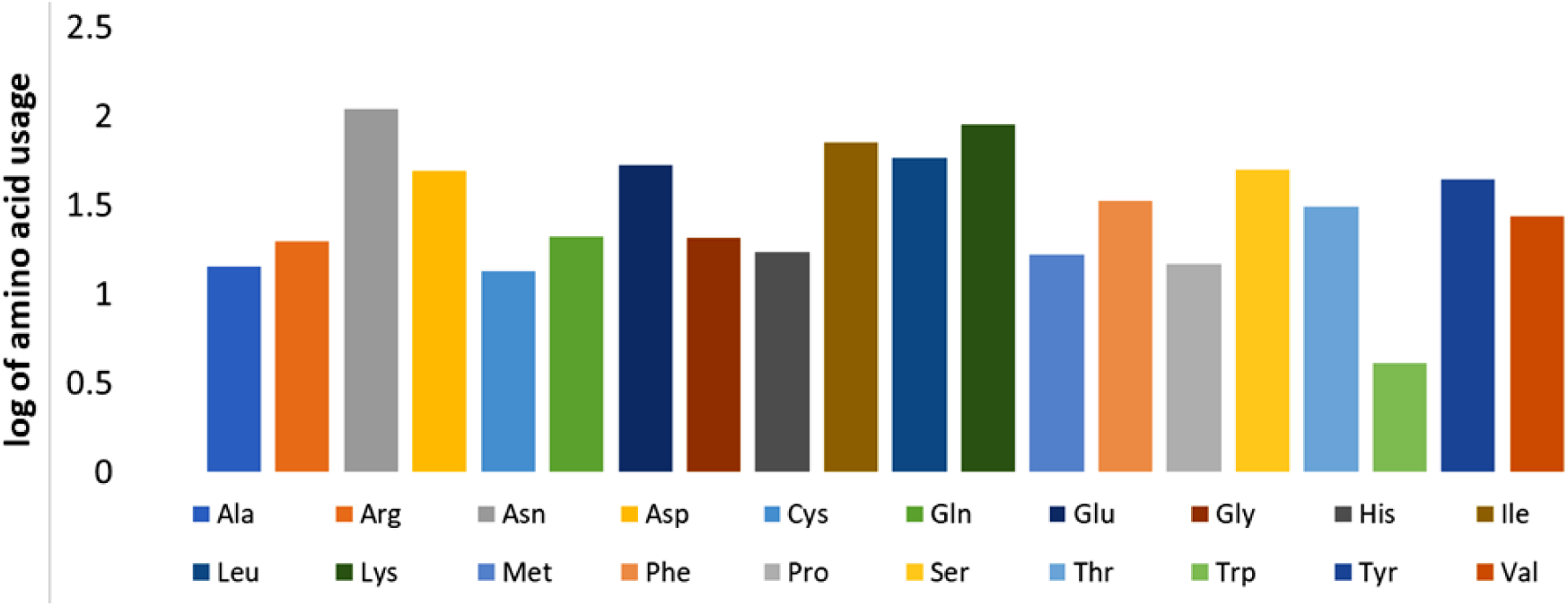
Amino acid usage among select *Plasmodium*

### Higher evolution of mitochondrial genomes suggests stronger pathogenicity

The Bayesian phylogenetic tree based on the mitochondrial genomes of select *Plasmodium* strains and some of their primary and secondary host grouped them in different clusters with considerable bootstrap value (Fig 3). The group consisting of all considered *Plasmodium* mitochondrial genomes was at one end of the tree. The group of secondary hosts was at another terminal whereas the group with a primary host was placed in between those two terminals connecting them with each other. This seems to be a representation of how the parasite has evolved its life to exploit both types of hosts (primary and secondary).

**Fig 3.**
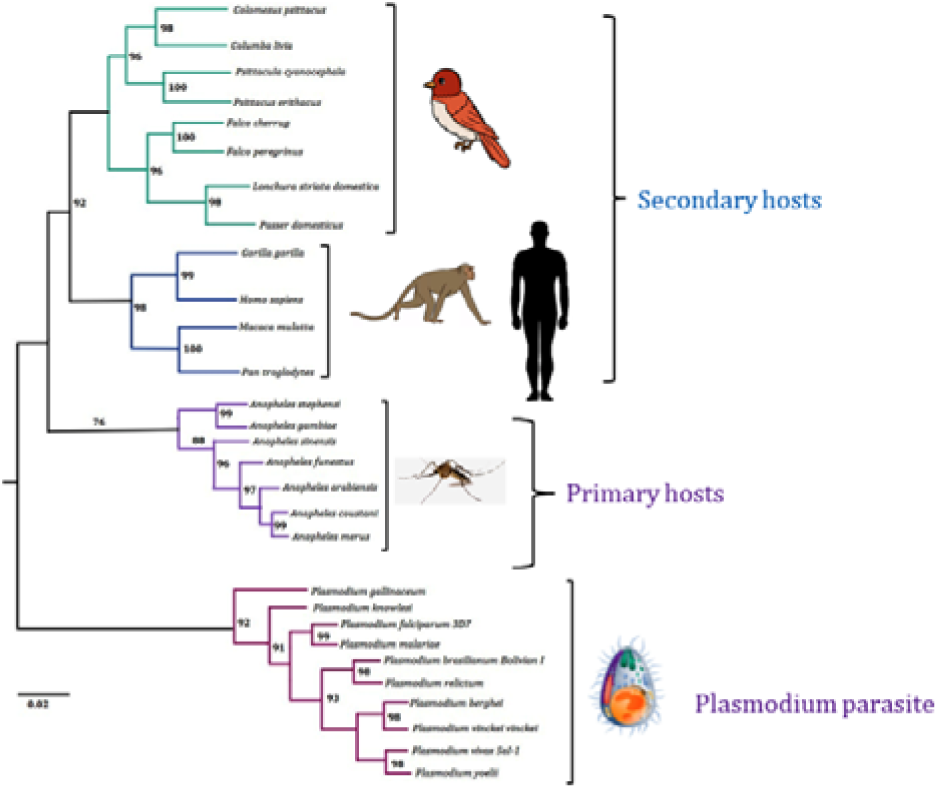
Bayesian phylogeny based on complete mitogenomes of select *Plasmodium* and their primary and secondary hosts.

A dN/dS based evolutionary study was performed on the select mitochondrial genomes and the result is represented in Fig 4. It was revealed that the evolutionary rate of the mitogenomes of *Plasmodium* was more (p<0.01) than both their primary and secondary host. This is a crucial point towards understanding the pathogenic behaviour of *Plasmodium*.

**Fig 4.**
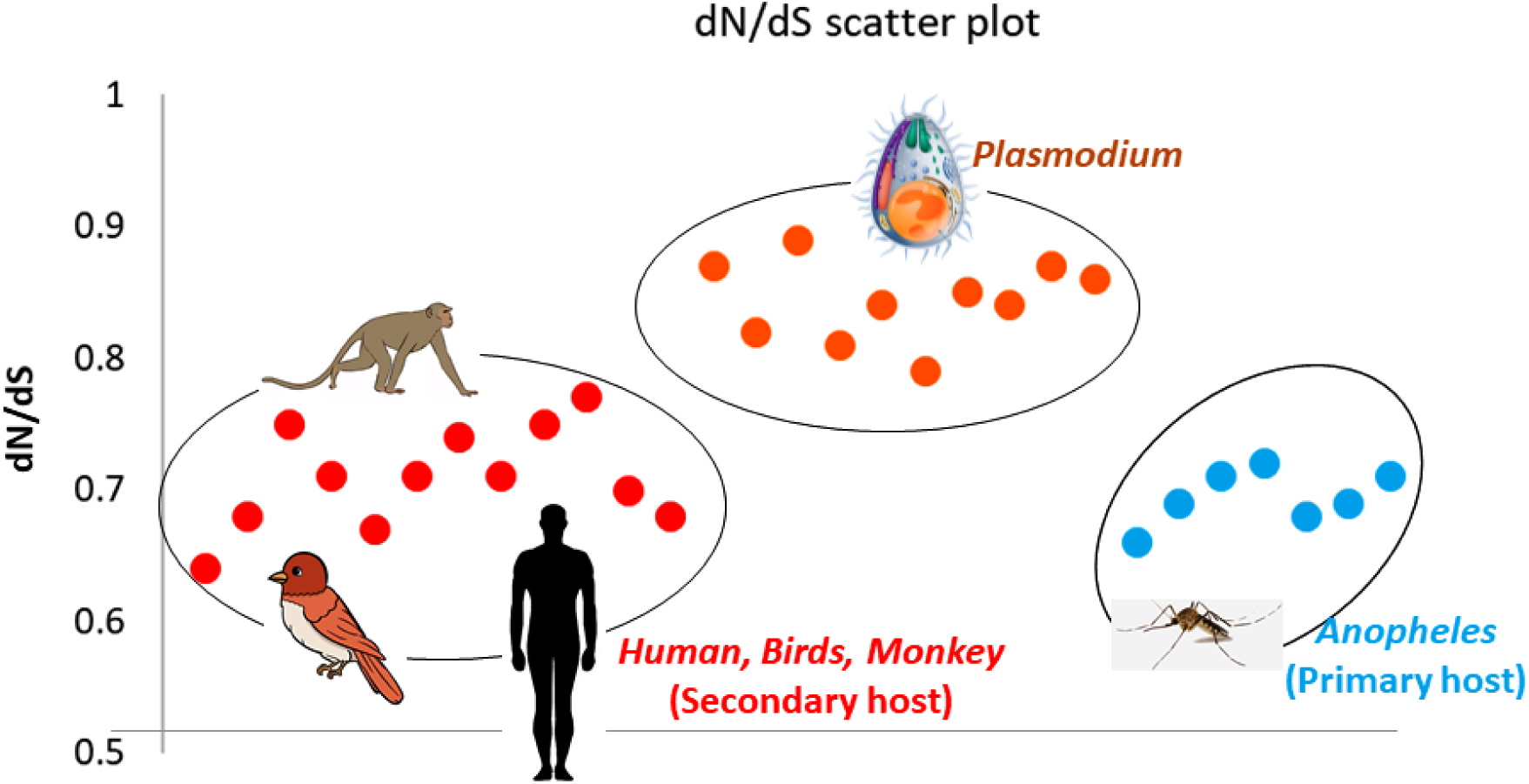
Differential evolutionary pattern between *Plasmodium* and their different hosts indicating rapid evolution of parasite mitogenome

### Understanding the host-parasite interaction through reverse ecology

The reverse ecology analysis was performed among select *Plasmodium* genomes and human (Homo sapiens) for a better understanding of the metabolic aspect of *Plasmodium* pathogenicity. The competition index (Fig 5) revealed that the inter-genus competition among *Plasmodium* is not very high (0.1-0.68). This indicates that a person can get affected by multiple *Plasmodium* strains at a time. Also, the competition index between *Plasmodium* and human was very high (0.8-0.97).

**Fig 5.**
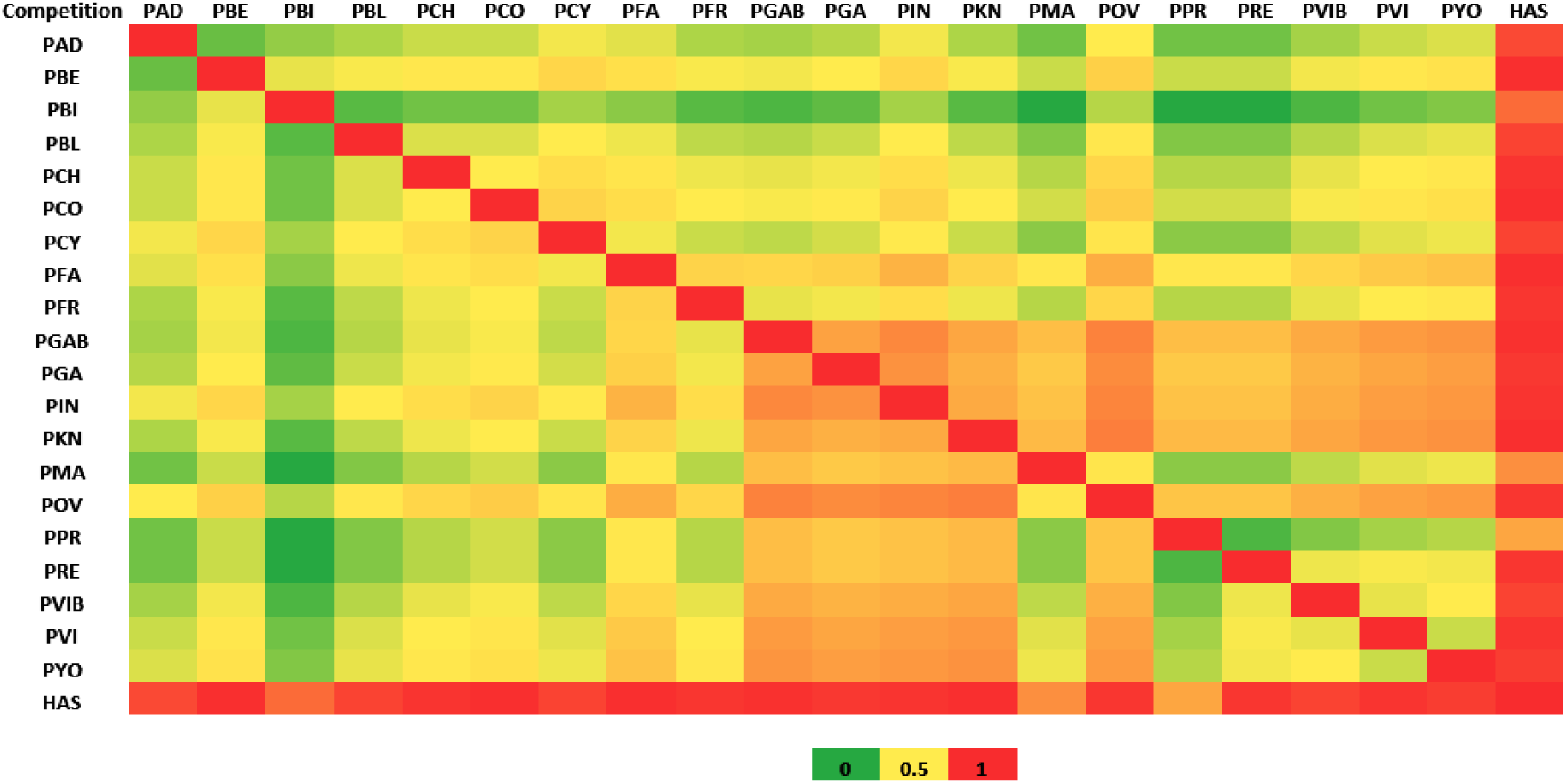
Heatmap on completion index among select *Plasmodium* and Homo sapiens. Color code has been indicated in the figure. The abbreviations are as follows: PAD=*Plasmodium adleri* G01, PBE=*Plasmodium berghei* ANKA, PBI=*Plasmodium billcollinsi* G01, PBL=*Plasmodium blacklocki* G01, PCH=*Plasmodium chabaudi* chabaudi, PCO=*Plasmodium coatneyi* Hackeri, PCY=*Plasmodium cynomolgi* strain B, PFA=*Plasmodium falciparum* 3D7, PFR=*Plasmodium fragile* strain nilgiri, PGAB=*Plasmodium gaboni* strain G01, PGA=*Plasmodium gallinaceum* 8A, PIN=*Plasmodium inui* San Antonio 1, PKN=*Plasmodium knowlesi* strain A1H1, PMA=*Plasmodium malariae* UG01, POV=*Plasmodium ovale* curtisi GH01, PPR=*Plasmodium praefalciparum* strain G01, PRE=*Plasmodium reichenowi* CDC, PVIB=*Plasmodium vinckei* brucechwatti DA, PVI=*Plasmodium vivax* P01, PYO=*Plasmodium yoelii* yoelii 17X, HSA=*Homo sapiens*

This clearly suggests that, once there is a malarial infection within the human body, the human immune system competes with this parasite. However, the human immune system can become weak since these parasites utilize the metabolic energy of the host itself to thrive within the human body. The competition between host and parasite is the main reason for all the symptoms of malaria. The complementation index (Fig 6) also supports the same hypothesis. The inter-genus complementation indicates malaria due to multiple *Plasmodium* strain infections. Also, the less complementation between humans and *Plasmodium* is obvious since both of them are competing with each other to thrive. This type of host-parasite interaction is rather complex and needs a strong biological interaction other than metabolic networking.

**Fig 6.**
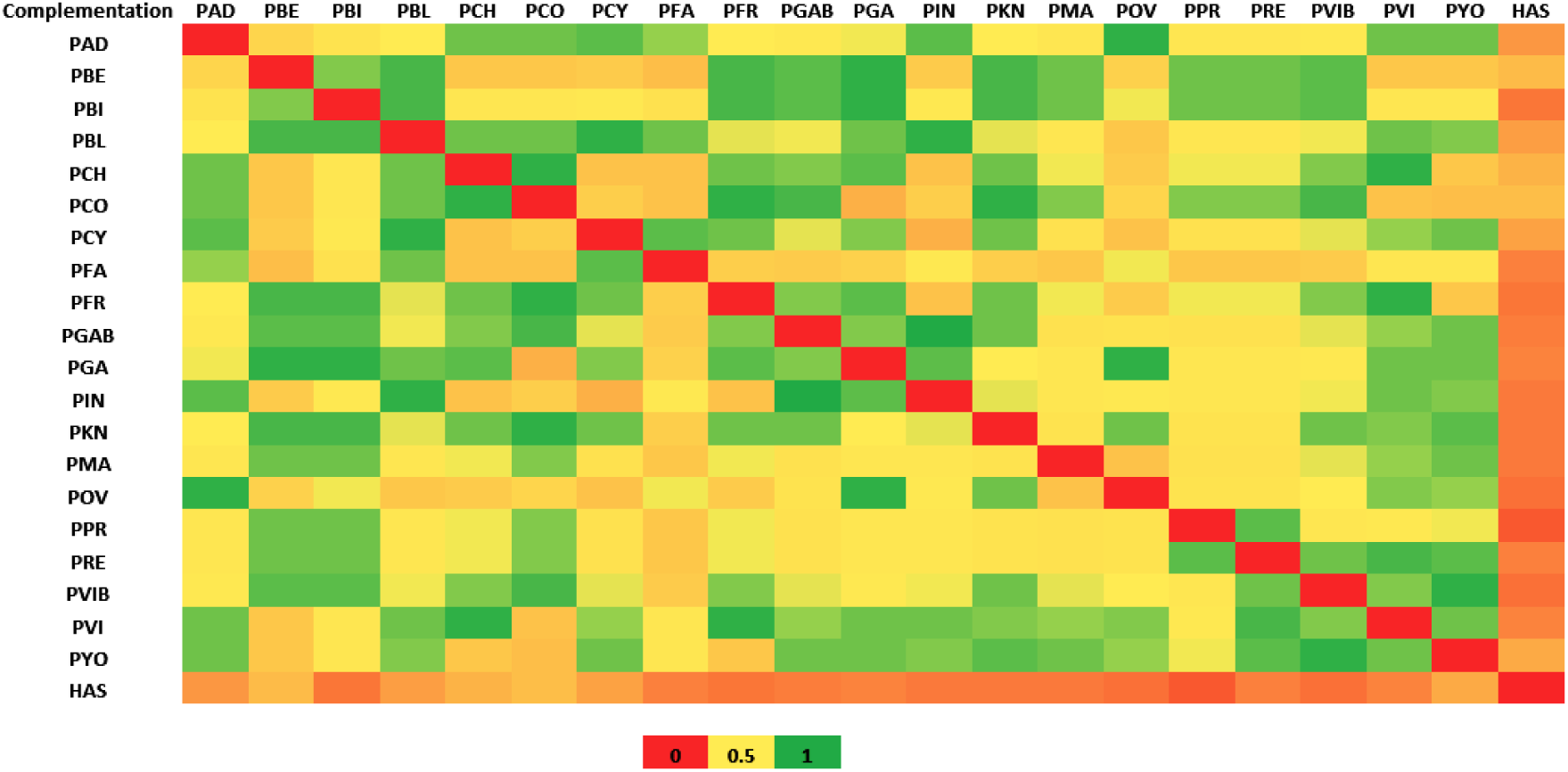
Heatmap on complementation index among select Plasmodium and Homo sapiens. Color code has been indicated in the figure. The abbreviations are as follows: PAD=*Plasmodium adleri* G01, PBE=*Plasmodium berghei* ANKA, PBI=*Plasmodium billcollinsi* G01, PBL=*Plasmodium blacklocki* G01, PCH=*Plasmodium chabaudi* chabaudi, PCO=*Plasmodium coatneyi* Hackeri, PCY=*Plasmodium cynomolgi* strain B, PFA=*Plasmodium falciparum* 3D7, PFR=*Plasmodium fragile* strain nilgiri, PGAB=*Plasmodium gaboni* strain G01, PGA=*Plasmodium gallinaceum* 8A, PIN=*Plasmodium inui* San Antonio 1, PKN=*Plasmodium knowlesi* strain A1H1, PMA=*Plasmodium malariae* UG01, POV=*Plasmodium ovale* curtisi GH01, PPR=*Plasmodium praefalciparum* strain G01, PRE=*Plasmodium reichenowi* CDC, PVIB=*Plasmodium vinckei* brucechwatti DA, PVI=*Plasmodium vivax* P01, PYO=*Plasmodium yoelii* yoelii 17X, HSA=*Homo sapiens*

### Pathogenicity of Plasmodium and host-parasite interaction

To explore more on the *Plasmodium* infection in humans, we exploited the system biology aspect. The set of proteins involved in malaria from both *Plasmodium* and humans were used in this case. It was found that *Plasmodium* utilizes twenty-four major proteins that cause malaria infection in humans (Supplementary Table 1). Those proteins mainly belong to erythrocyte membrane-binding protein, merozoite surface protein, circumsporozoite (CS) protein, reticulocyte binding protein, rifin, stevor, perforin-like protein, cell traversal protein, apical membrane antigen, thrombospondin-related anonymous protein and sporozoite protein essential for cell traversal. The biological interaction among them was very strong. The protein-protein interaction (PPI) enrichment p-value was < 1.0e^-16^. The average local clustering coefficient was 0.758 (Fig 7). The gene ontology (GO) and enrichment study revealed all of those 24 proteins were associated with malaria pathogenicity.

**Fig 7.**
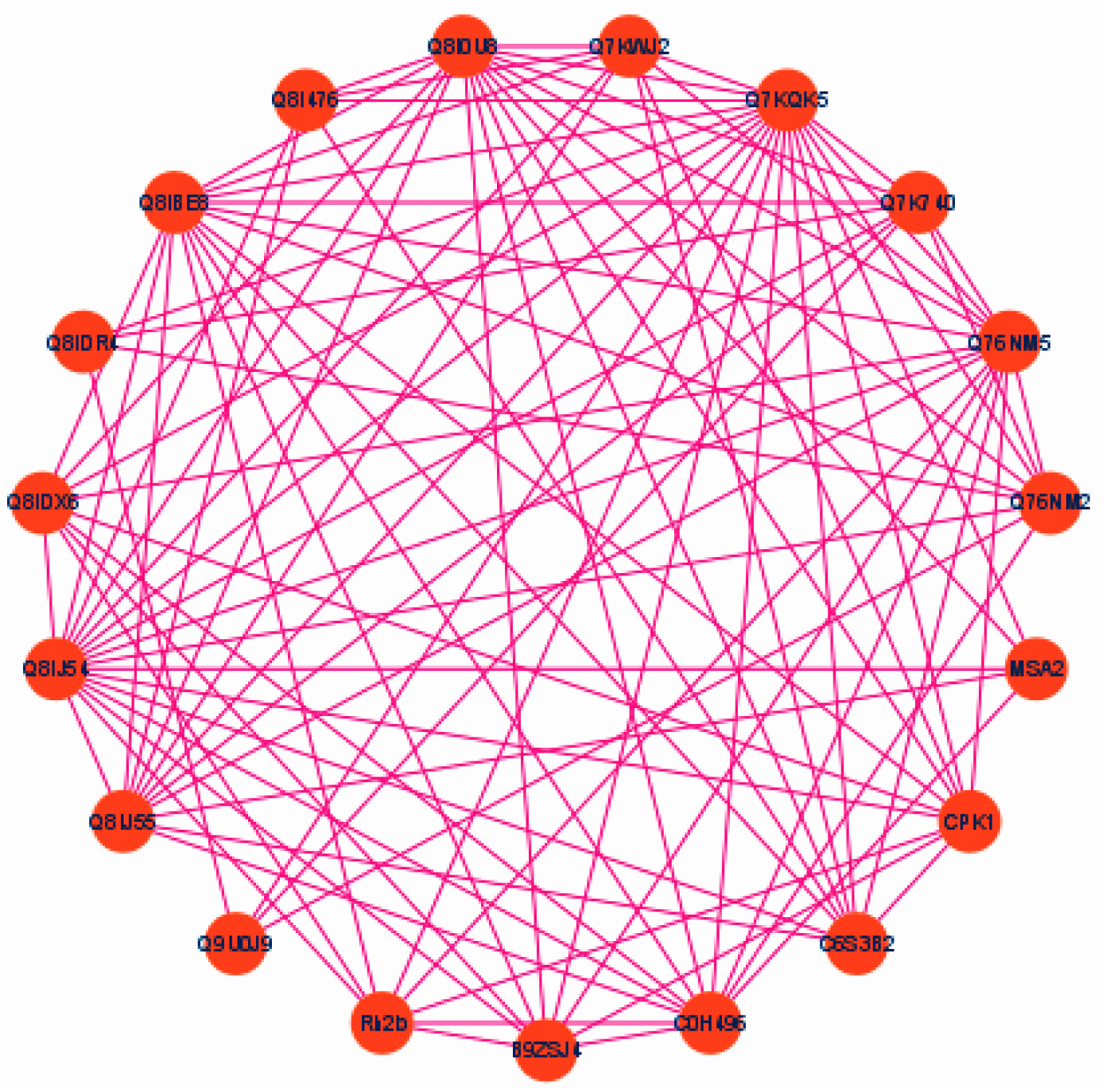
Biological network among the virulence proteins in *Plasmodium*

On other hand, forty-nine different proteins were found to be related to malaria in humans. Those proteins were majorly from selectin, glycophorin, haemoglobin subunit alpha, and the erythrocyte membrane protein family. The PPI enrichment p-value was< 1.0e^-18^ with an average local clustering co-efficient of 0.851 (Fig 8).

**Fig 8.**
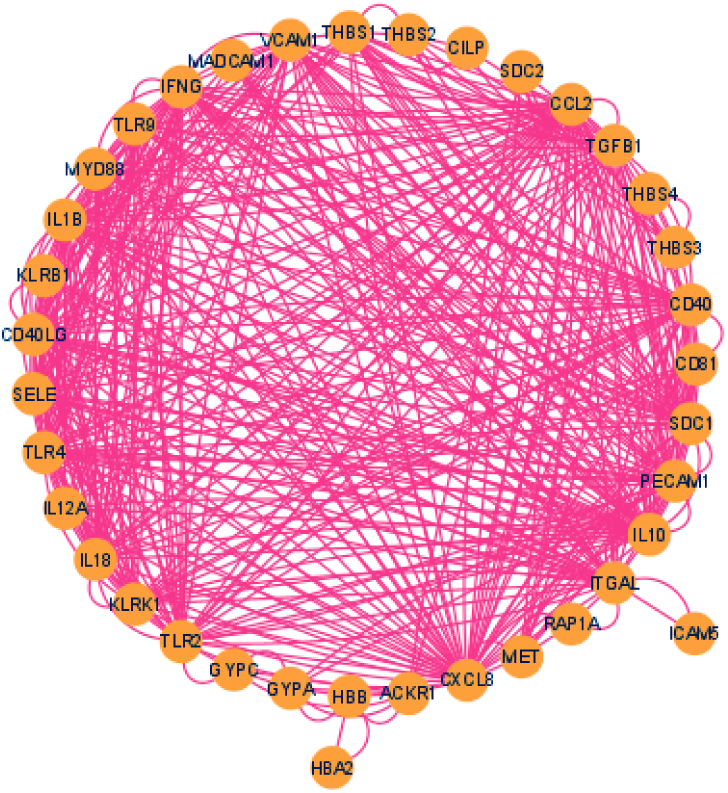
Biological network among the Malaria related proteins in Human

The GO and enrichment analysis revealed that these proteins were broadly associated with immunity, B-cell and T-cell maturation, natural killer cell mediated cytotoxicity, positive regulation of matrix metallopeptidase secretion, and regulation of interleukins along with positive regulation of granulocyte-macrophage colony-stimulating factor production.

Another biological network analysis was performed between the *Plasmodium* and human and the PPI enrichment p-value of that network was < 1.0e^-16^ with an average local clustering co-efficient of 0.721. It was found that selectin, glycophorin, haemoglobin subunit alpha, and erythrocyte membrane protein families of humans interacted with all the major *Plasmodium* proteins to win over this parasite (Fig 9).

**Fig 9.**
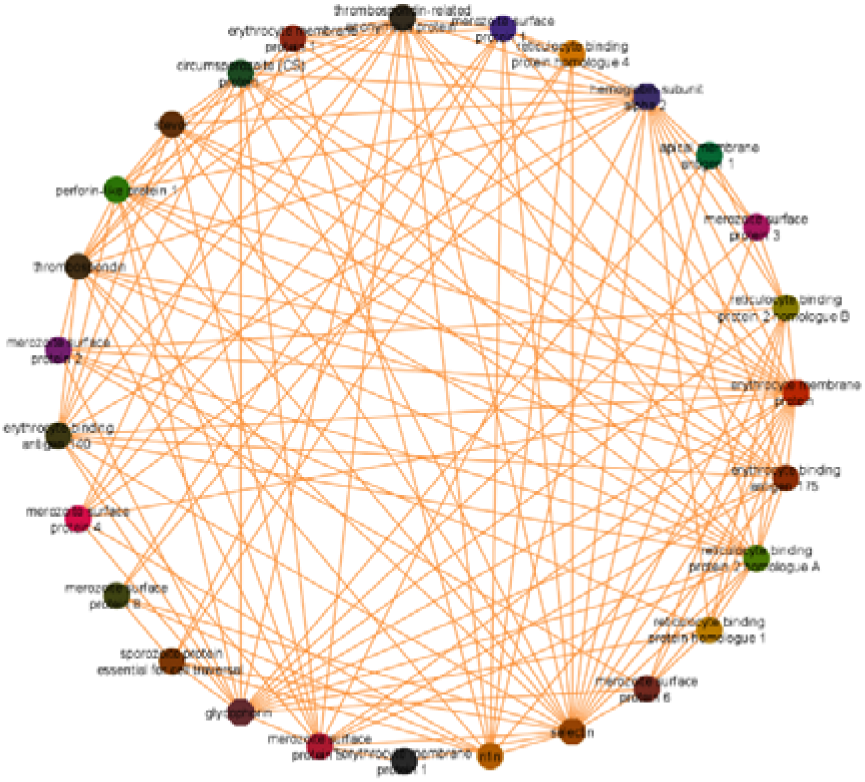
Biological network among Malaria specific protein of *Plasmodium* and Human

### Multi-drug resistance of *Plasmodium*

The ABC transporter protein, efflux pumps, and multi-drug efflux pumps are the proteins mainly associated with the emergence of the drug resistant *Plasmodium* strains.

The biological network analysis gave a PPI enrichment p-value < 1.0e^-18^ with an average local clustering co-efficient 0.698. GO enrichment analysis revealed ATPase-coupled transmembrane transporter activity, transmembrane transporter activity, ATPase activity, ATP binding and catalytic activity were major biological processes associated with those proteins. This strong interaction among these proteins is the major reason the drugs against *Plasmodium are* failing to prevent or cure malaria (Fig 10).

**Fig 10.**
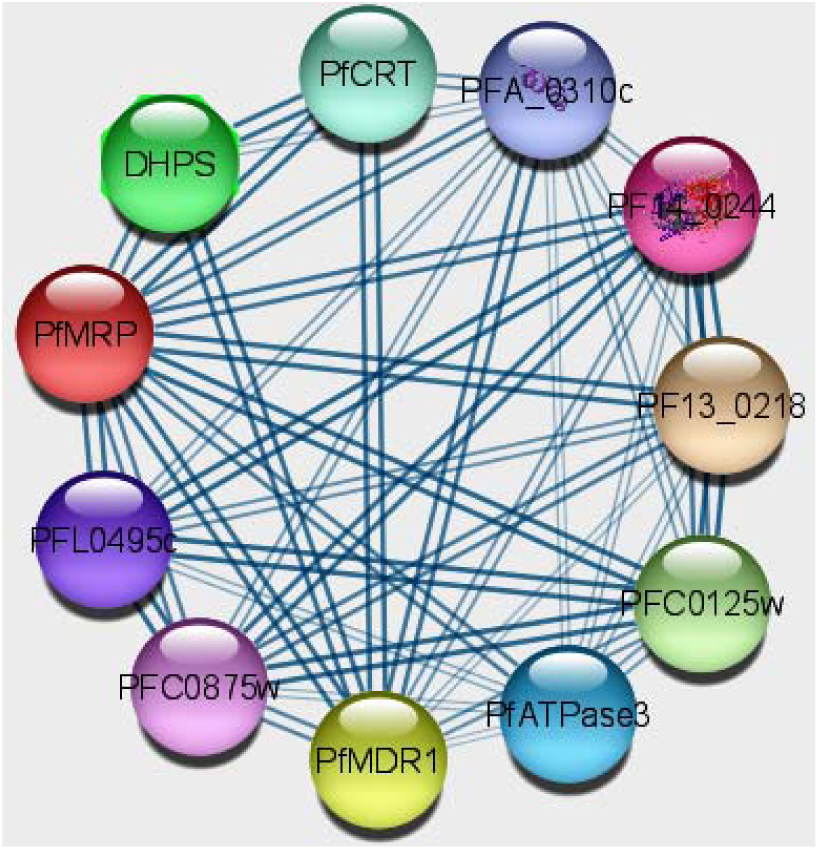
Biological network among *Plasmodium* proteins related their drug resistance property

### Cymbopogon against the Plasmodium and mechanism of action

Thirty-five phyto-compounds derived from *Cymbopogon* sp. were docked with the select target proteins (PDB id: 2BL9, 6VTY) (Table 2).

Cymbopogonol, Triterpenoid, Cymbopogone, Phytosterols, Beta-caryophyllene, luteolin, Sesquiterpene II alcohol diglucoside (Compound 1), Homoorientin and p-Camphorene were found to bind with both target protein with binding energy more than -7.0kcal/mol and were considered for further analysis (Table 2, Fig 11, Supplementary file 1). All those compounds obeyed Lipinski’s rule and were passed into PAINS and Veber indicating each of them can be used as drug molecules in humans without much concern. Among those compounds, Cymbopogonol was found to be the best compound against both 2BL9 and 6VTY with dock score of -15.1 kcal/mol and -15.7kcal/mol respectively (Fig 12, Fig 13).

**Fig 11.**
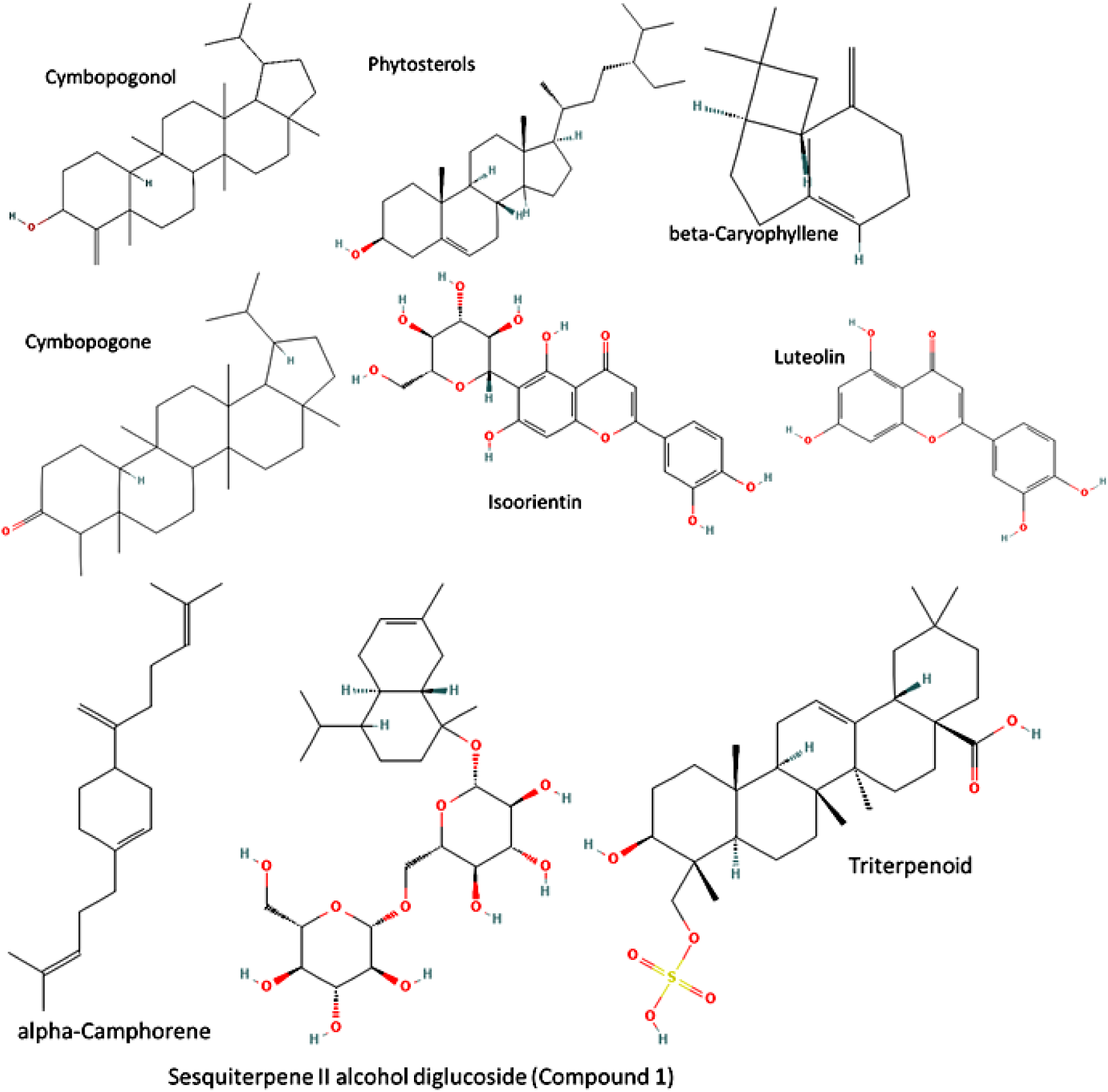
Phytocompound structures showed dock score more than -7.0 Kcal/mol with both 2BL9 and 6VTY

**Fig 12.**
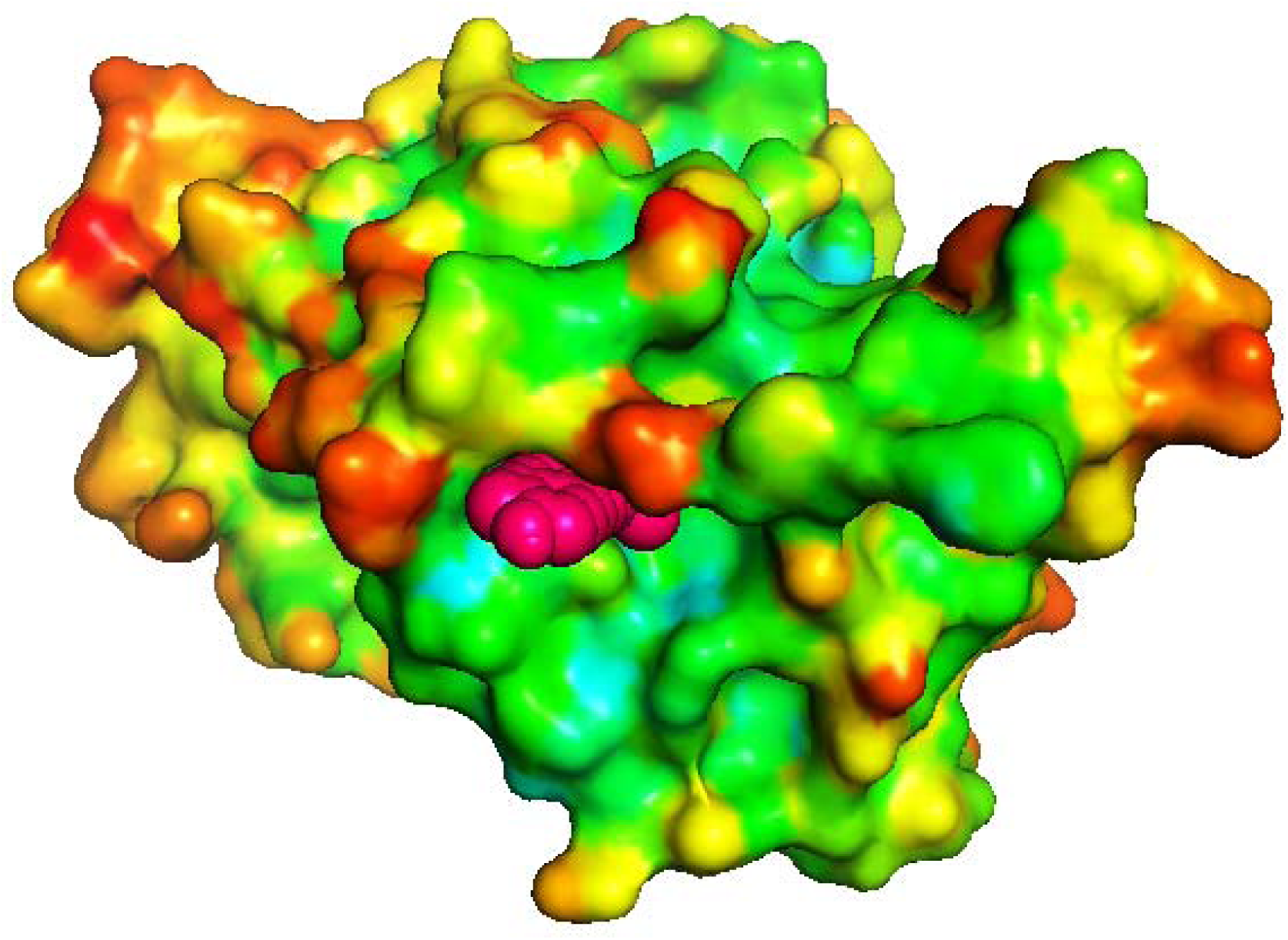
The binding of Cymbopogonol with 2BL9

**Fig 13.**
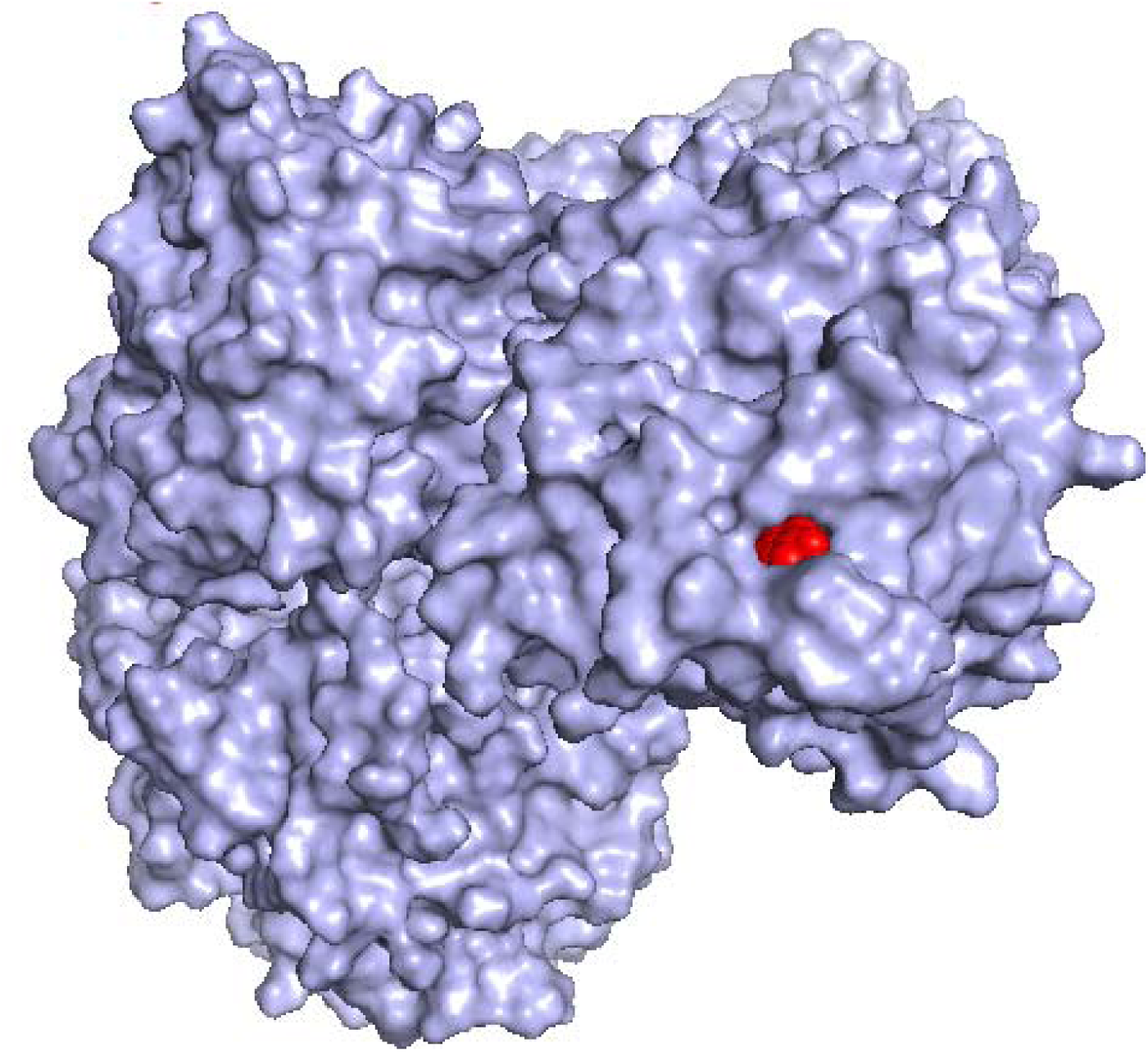
The binding of Cymbopogonol with 6VTY

The interacting amino acids between the Cymbopogonol-2BL9 complex were 191Leu, 193 Phe, 209 Phe, 211Glu and 235Tyr. This site was not the same as the site where pyrimethamine binds in the crystal structure 2BL9. Similarly, the interacting site of Cymbopogonol-6VTY was composed of 171Phe, 172Leu, 176 Leu, 185His, 188Phe, 191Leu and 531Val. This site was also different from the site where the pyrrole-based inhibitor was bound in the crystal structure of 6VTY. Since *Plasmodium* is now becoming resistant to all known drugs these new interacting sites and *Cymbopogon* sp. derived phyto-compounds may prove to be beneficial against the drug-resistant strains.

Both Cymbopogonol-2BL9 and Cymbopogonol-6VTY were further used for molecular dynamics and simulation study. The 100ns simulation study revealed no significant change between the apo-protein and the receptor-protein complex. The RMSD plot (Fig 14) of Cymbopogonol-2BL9 showed the fluctuation of the carbon backbone (of the complex) was reduced after 50ns and was stabilized by 100ns (green colour). Severe fluctuation of Cymbopogonol-6VTY carbon backbone was absent throughout the 100ns time and it was further reduced at 99ns time (Fig 15).

**Fig 14.**
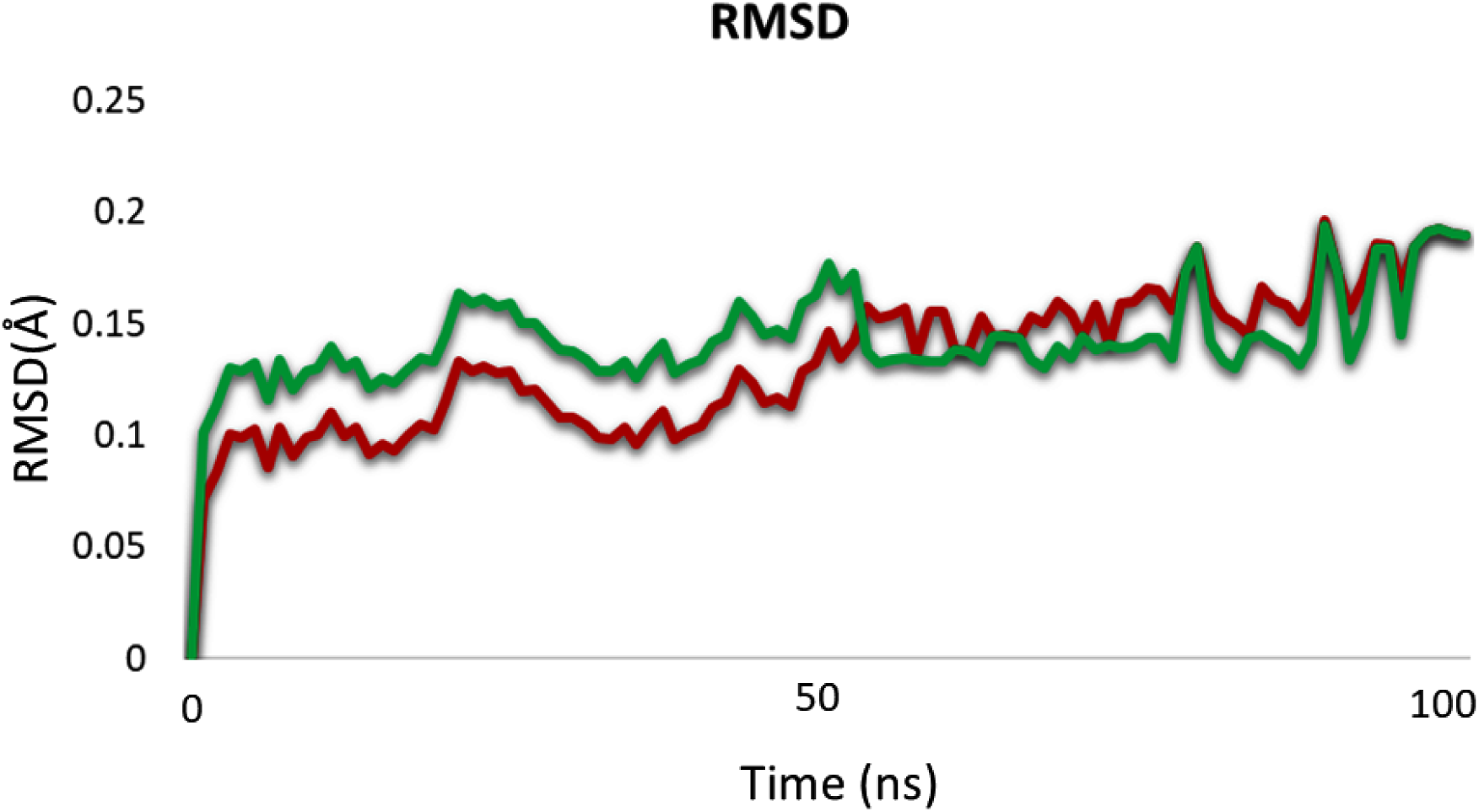
RMSD plot for Cymbopogonol-2BL9 complex

**Fig 15.**
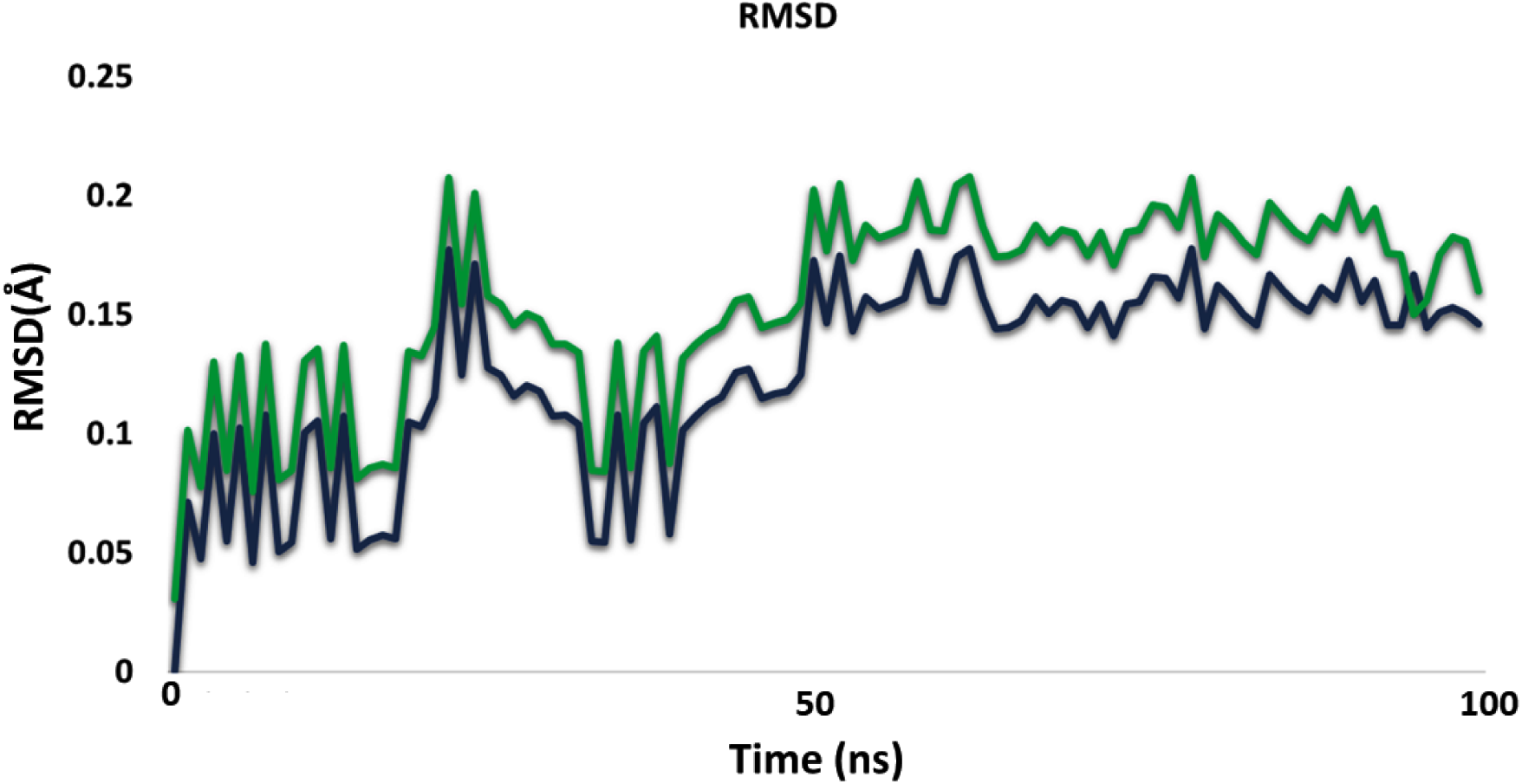
RMSD plot for Cymbopogonol-6VTY complex

Both the RMSF plots (Fig 16, Fig17) also revealed the stabilization of the carbon backbone of both Cymbopogonol-2BL9 and Cymbopogonol-6VTY within a 100ns time scale. The MM-GBSA calculation revealed the free energies to be-16.25 kcal/mol and -15.89kcal/mol for Cymbopogonol-2BL9 and Cymbopogonol-6VTY respectively.

**Fig 16.**
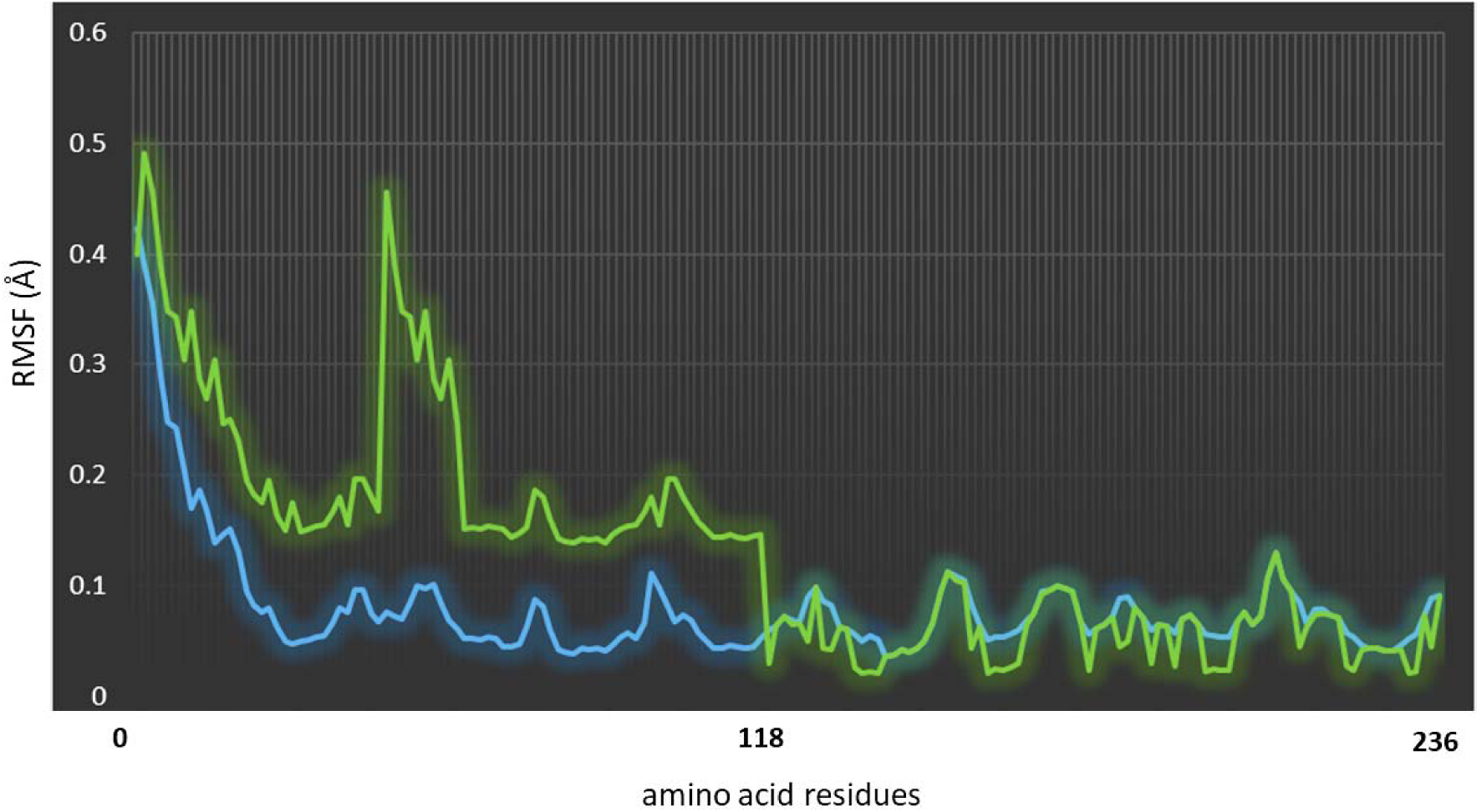
RMSF plot for Cymbopogonol-2BL9 complex

**Fig 17.**
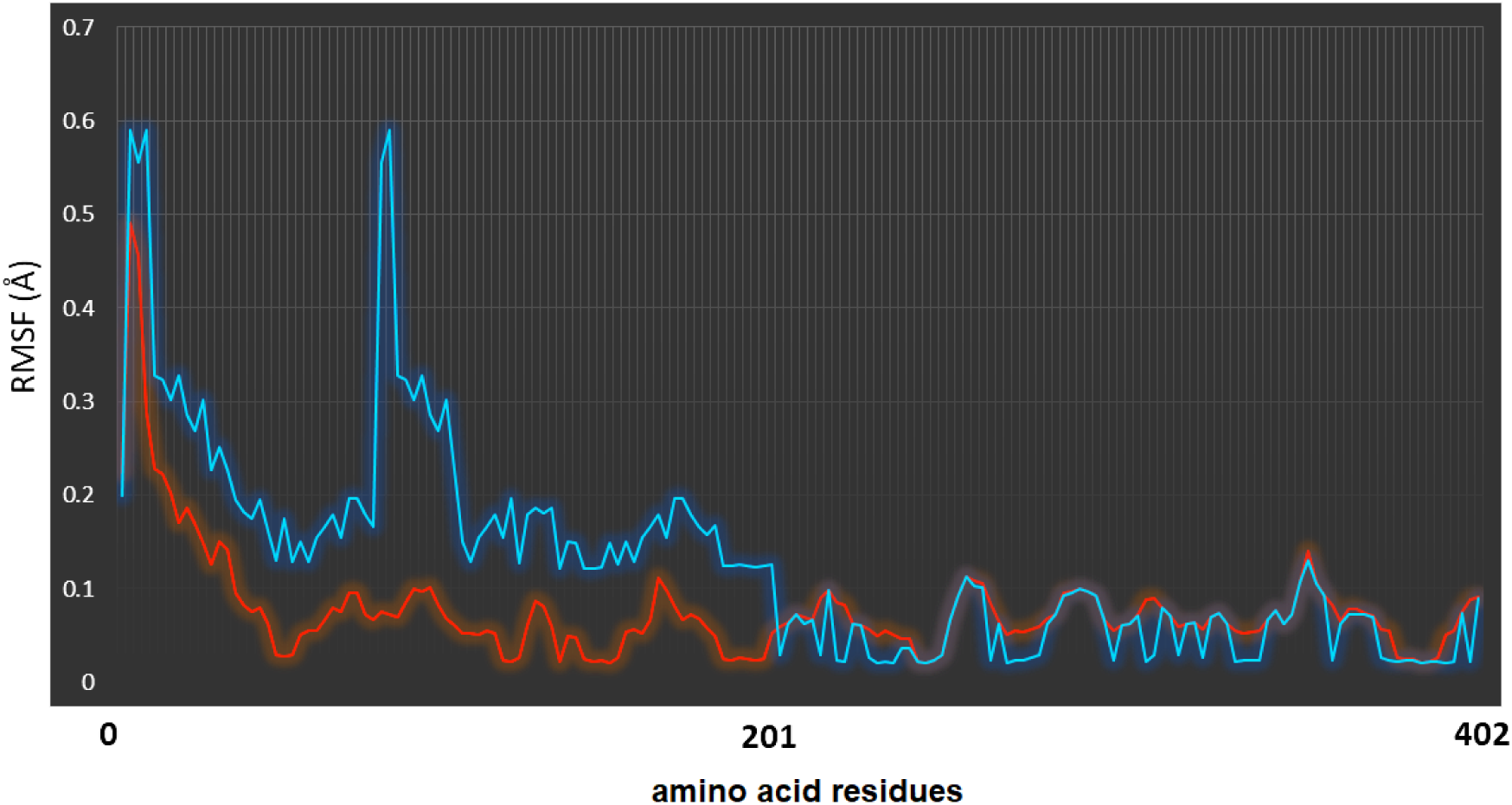
RMSF plot for Cymbopogonol-6VTY complex

## Discussion

### AT-rich compositional constrain mediated translational efficiency persists among Plasmodium

The *Plasmodium* genomes and mitogenomes were found to be highly AT-rich. This result was supported by previous findings (Musto et al. 1997; Musto et al. 1999). Synonymous codon usage is not a random phenomenon. The ‘genomic hypotheses’ has proposed that codon bias is generally organism-specific (Musto et al. 1999). The high AT preference among *Plasmodium* is a direct proof of their compositional constrain. Moreover, the AT biased nature of PHX was more than PLX. This can be contributed by two factors – (a) compositional constrain and (b) natural selection (Musto et al. 1999). The correlation among other codon usage indices like Fop, CAI, ITE and Enc with AT and AT3 indicated translational efficiency among the select *Plasmodium* strains. The Enc was negatively correlated with AT revealing effect of the natural selection on those strains. ITE was positively correlated with AT, AT3, CAI and Fop indicating efficient translational machinery among *Plasmodium*.

### Codon usage pattern of Plasmodium points towards host-adaptivity

Previous studies have revealed the connection between codon adaptation and ecological preference (Peden, 2000). There is also a relation between codon evolution and codon adaptation. The synonymous codon usage pattern of *Plasmodium* was compared with humans and *Anopheles*. The twenty-two AT-rich codons were found to be preferred among all select *Plasmodium* and their primary and secondary hosts. This indicated a codon co-evolution and co-adaptivity. Since most of those aforementioned codons were AT-rich and transitional efficiency among *Plasmodium* is AT driven, we may propose that this codon co-evolution is helping the parasite to live within the host systems as well as to utilize their metabolic resources (Botzman and Margalit, 2011). Thus, codon usage might be playing a crucial role in the host-parasite interaction strategy. Moreover, the amino acids preferred by *Plasmodium* are not energy-costly aromatic amino acids rather they are energy-economic. Proteins needed for the co-evolution and co-adaptivity with host are mostly made of energy economic amino acids among *Plasmodium* and are thus can be translated without excess energy expenditure. This again indicated sophisticated translational machinery among *Plasmodium*.

### Rapid evolution of Parasites supports their pathogenicity

Rapid evolution in parasites maximizing their reproductive success and transmission potential (either from host to host or from the environment to host) ultimately results in uncontrolled parasitic infection among hosts. During the host-parasite co-evolution, reciprocal genetic changes happen in both the host and parasites. Parasitic virulence is mostly the effect of such dynamic genetic changes which often reduce the fitness level of hosts (Rafaluk et al. 2015). The optimal virulence can be reached by an evolutionary compromise between the too rapid and too slow reproduction rate of parasites within the host body. Too rapid reproduction can affect the transmission rate of parasites from one host to another since it may kill the host itself even before the transmitting stage of the parasites is reached (Rafaluk et al. 2015). This is like a trade-off between transmission and virulence which is based on the simple idea that an increased transmission rate costs the duration of the infection cycle. On the other hand, with too slow reproduction of parasites, it would not be possible to exploit the host resources at a maximum level. Thus, a complex regulatory mechanism exists among the genetic diversity, reproduction rate, transmission rate and evolution of parasites. In this study, we compared the mitogenomic evolution of *Plasmodium*, *Anopheles* (primary host) and human (secondary host) and found that a rapid evolution rate persists among *Plasmodium* mitogenomes. The higher evolution rate of *Plasmodium* mitogenomes in comparison to their hosts is a direct strategy to combat the host defence mechanism. This rapid evolution rate may also aid in the emergence of drug-resistance strains of *Plasmodium*. With the errors acquired during genomic replication and the lack of endogenous repair capacity in mitogenomes, the mutational rate of mtDNA remains exceptionally higher than nuclear DNA (Jones et al. 2021). Mutation(s) resulting in functional variants related to maternal lineages are clinically important and are also associated with drug susceptibility or resistance. Hence the rapid evolution of *Plasmodium* mitogenomes is indicating the pivotal role of mitogenomes in the emergence of drug-resistant strains of *Plasmodium*.

### PPI analysis indicates complex Plasmodium -Human interaction

The reverse ecology analysis proved a strong competition between *Plasmodium* and human. Moreover, strong biological interactions were found both in *Plasmodium* and human among the proteins associated with malaria indicating a complex host-parasite interaction. It was clear from the PPI analysis that, the main target of *Plasmodium* is the human immune system. *Plasmodium takes* different strategies at different parts of their life cycle to invade the line of defence. Free sporozoites and intrahepatic parasites have to bypass the hurdle of host immune response before they reach the erythrocytic stage (Belachew, 2018). Sporozoites can pass through the Kupffer cells and some endothelial cells actively. Some of the sporozoites use the gap between those cells to avoid this line of defence. This is very intriguing. The Kupffer cells are well-known for their cell-killing nature however, the *Plasmodium* sporozoites have no problem with them and that’s due to circumsporozoite protein (CSP) (Belachew, 2018). CSP binds to Kupffer cells’ surface proteins producing high level of intracellular cAMP/EPAC which prevents the ROS (reactive oxygen species) formation (26). The interaction between CSP and Kupffer cells also downregulates the Th1 cytokines (inflammatory) and up-regulates Th2 cytokines (anti-inflammatory) (Belachew, 2018). In some cases, the CSP binding causes Kupffer cells apoptosis and reduces major histocompatibility complex (MHC)-I expression level. Thus, sporozoites with the help of CSP can manipulate the Kupffer cells and enter into the erythrocytic stage (Belachew, 2018). Lack of MHC-I helps the parasites to avoid the CD8^+^ T cells mediated killing. Two proteins Rifins and Stevors are important for immune evasion at the mature trophozoite stage (Belachew, 2018). Thus, *Plasmodium* proteins can modulate the human immune system and can bypass its attack.

The antigenic variation of *Plasmodium* due to Rifin, Stevor and other related proteins are major challenges to producing an anti-malaria vaccine (Belachew, 2018). Although several drugs have been discovered and were proved to be effective against malaria, the recent emergence of drug-resistant *Plasmodium* strains are creating recurrent concerns. The strong biological interaction among ABC transporters, efflux pumps and drug transporters indicated towards complex and advanced machinery of drug resistance property of *Plasmodium*.

### Proposal of Cymbopogonol as a novel antimalarial drug

Failure of existing drugs to combat *Plasmodium* is pointing toward the need for new drug targets and new drug molecules. We have already mentioned the potency of mitogenome coding proteins as drug targets (material and method section). From molecular docking analysis, we found Cymbopogonol was the best-docked compound and we have identified new targeting sites from both 2BL9 and 6VTY protein. Inhibition of 2BL9 (dihydrofolate reductase) and 6VTY (Dihydroorotate dehydrogenase) by Cymbopogonol can directly disturb the mitochondrial functionality and thus cease the growth of *Plasmodium* in human. The physiochemical properties along with molecular dynamics results also support the credibility of Cymbopogonol as an anti-*Plasmodium*.

## Conclusion

Malaria is still a leading cause of death worldwide. The causative agent, *Plasmodium* has emerged as a strong multi-drug resistance organism. We are in immense need of new therapeutic strategies against *Plasmodium* and for that, we need a better understanding of the *Plasmodium* genomes, their evolution and host-adaptive nature. In this study, we considered 55 *Plasmodium* genomes and studied their codon and amino acid usage, evolutionary pattern, and interaction strategies with the host. From the results, it was revealed that AT compositional constrain drive the translational efficacy among *Plasmodium*. The shared pattern of codon usage among *Plasmodium*, *Anopheles* and humans directly represented the co-evolution aspect of host-parasite interaction. The rapid evolution of *Plasmodium* mitogenomes supported the optimal virulence and transmission rate of *Plasmodium* in both primary and secondary hosts. The reverse ecology analysis indicated a strong competition between *Plasmodium* and humans regarding resource utilization. Protein-protein interaction (PPI) enrichment study suggested a strong interaction between the virulent proteins of *Plasmodium* and the human immune system. Finally, two new drug targeting sites were found from *Plasmodium* mitogenome coding proteins and Cymbopogonol derived from Lemongrass was proposed as a novel drug molecule against *Plasmodium.* This is the first study reporting Cymbopogonol as a possible drug candidate against *Plasmodium* and is open for clinical trials.

### Conflict of Interest

The authors declare no conflict of interest.

### Authors Contribution

RPS conceived the idea. IS, SSR and GDV performed all the analyses. RPS, IS, SSR and GDV wrote the manuscript. All authors have agreed with the final version of the manuscript.

## Acknowledgements

IS acknowledges SACON Coimbatore for providing the facilities needed for this study. SSR acknowledges Mahatma Gandhi Central University. GDV acknowledges Bihar Veterinary College and RPS acknowledges Central University of South Bihar.

## Notes

### Competing Interest Statement

The authors have declared no competing interest.

## Reference

Achan, J., Serwanga, A., Wanzira, H., Kyagulanyi, T., Nuwa, A., Magumba, G., Kusasira, S., Sewanyana, I., Tetteh, K., Drakeley, C. and Nakwagala, F., 2022. Current malaria infection, previous malaria exposure, and clinical profiles and outcomes of COVID-19 in a setting of high malaria transmission: an exploratory cohort study in Uganda. Lancet Microbe, 3(1), pp.e62–e71.

Arnot, D.E. and Gull, K., 1998. The Plasmodium cell-cycle: facts and questions. Ann. trop. med. parasitol., 92(4), pp.361–365.

Belachew, E.B., 2018. Immune response and evasion mechanisms of Plasmodium falciparum parasites. J. Immunol. Res.. https://doi.org/10.1155/2018/6529681

Botzman, M. and Margalit, H., 2011. Variation in global codon usage bias among prokaryotic organisms is associated with their lifestyles. Genome Biol., 12(10), pp.1–11.

Bryant, C., Voller, A. and Smith, M.J.H., 1964. The Incorporation of Radioactivity from [14C] Glucose into the Soluble Metabolic Intermediates of Malaria Parasites. Am J Trop Med Hyg, 13(4), pp.515–19.

Chukwuocha, U.M., Fernández-Rivera, O. and Legorreta-Herrera, M., 2016. Exploring the antimalarial potential of whole Cymbopogon citratus plant therapy. J. Ethnopharmacol., 193, pp.517–523.

Eid, M.M., 2021. Co-infection with COVID-19 and Malaria in a Young Man. Dubai Med. J., 4(2), pp.164–166.

Fleck, S.L., Pudney, M. and Sinden, R.E., 1996. The effect of atovaquone (566C80) on the maturation and viability of Plasmodium falciparum gametocytes in vitro. Trans. R. Soc. Trop. Med. Hyg., 90(3), pp.309–312.

Galinski, M.R., Barnwell, J.W., Abee, C.R., Mansfield, K., Tardif, S. and Morris, T., 2012. Nonhuman primate models for human malaria research. Nonhuman primates in biomedical research. Editors: Christian Abee, Keith Mansfield, Suzette Tardif, Timothy Morris, Hardcover ISBN: 9780123813657, eBook ISBN: 9780123978370

Haldar, K., Bhattacharjee, S. and Safeukui, I., 2018. Drug resistance in Plasmodium. Nat. Rev. Microbiol., 16(3), pp.156–170.

Handunnetti, S.M., Mendis, K.N. and David, P.H., 1987. Antigenic variation of cloned Plasmodium fragile in its natural host Macaca sinica. Sequential appearance of successive variant antigenic types. J. Exp. Med., 165(5), pp.1269–1283.

Jones, S.W., Ball, A.L., Chadwick, A.E. and Alfirevic, A., 2021. The Role of Mitochondrial DNA Variation in Drug Response: A Systematic Review. Front. Genet., p.1430.

Ke, H., Morrisey, J.M., Ganesan, S.M., Painter, H.J., Mather, M.W. and Vaidya, A.B., 2011. Variation among Plasmodium falciparum strains in their reliance on mitochondrial electron transport chain function. Eukaryot. Cell, 10(8), pp.1053–1061.

Kokkonda, S., Deng, X., White, K.L., El Mazouni, F., White, J., Shackleford, D.M., Katneni, K., Chiu, F.C., Barker, H., McLaren, J. and Crighton, E., 2020. Lead optimization of a pyrrole-based dihydroorotate dehydrogenase inhibitor series for the treatment of malaria. J. Med. Chem., 63(9), pp.4929–4956.

Kongsaeree, P., Khongsuk, P., Leartsakulpanich, U., Chitnumsub, P., Tarnchompoo, B., Walkinshaw, M.D. and Yuthavong, Y., 2005. Crystal structure of dihydrofolate reductase from Plasmodium vivax: pyrimethamine displacement linked with mutation-induced resistance. Proc. Natl. Acad. Sci., 102(37), pp.13046–13051.

Lunev, S., Batista, F.A., Bosch, S.S., Wrenger, C. and Groves, M.R., 2016. Identification and validation of novel drug targets for the treatment of Plasmodium falciparum malaria: new insights. InTech. Edited by Alfonso J. Rodriguez-Morales, DOI: 10.5772/65659

Maguire, S.E., Afify, A., Goff, L.A. and Potter, C.J., 2022. Odorant-receptor-mediated regulation of chemosensory gene expression in the malaria mosquito Anopheles gambiae. Cell Rep., 38(10), p.110494.

Moreno, A., Cabrera-Mora, M., Garcia, A., Orkin, J., Strobert, E., Barnwell, J.W. and Galinski, M.R., 2013. Plasmodium coatneyi in rhesus macaques replicates the multisystemic dysfunction of severe malaria in humans. Infect. Immun., 81(6), pp.1889–1904.

Musto, H., Cacciò, S., Rodríguez-Maseda, H. and Bernardi, G., 1997. Compositional constraints in the extremely GC-poor genome of Plasmodium falciparum. Mem. Inst. Oswaldo Cruz., 92, pp.835–841.

Musto, H., Romero, H., Zavala, A., Jabbari, K. and Bernardi, G., 1999. Synonymous codon choices in the extremely GC-poor genome of Plasmodium falciparum: compositional constraints and translational selection. J. Mol. Evol., 49(1), pp.27–35.

Ngoubangoye, B., Boundenga, L., Arnathau, C., Mombo, I.M., Durand, P., Tsoumbou, T.A., Otoro, B.V., Sana, R., Okouga, A.P., Moukodoum, N. and Willaume, E., 2016. The host specificity of ape malaria parasites can be broken in confined environments. Int. J. Parasitol., 46(11), pp.737–744.

Peden, J.F., 2000. Analysis of codon usage. Thesis submitted to the University of Nottingham for the Degree of Doctor of Philosophy, August, 1999

Picot, S., Bienvenu,AL. 2020, Plasmodium, Reference Module in Biomedical Sciences, Elsevier, ISBN 9780128012383, https://doi.org/10.1016/B978-0-12-818731-9.00041-0.

Planche, T. and Krishna, S., 2006. Severe malaria: metabolic complications. Curr. Mol. Med., 6(2), pp.141–153.

Rafaluk, C., Gildenhard, M., Mitschke, A., Telschow, A., Schulenburg, H. and Joop, G., 2015. Rapid evolution of virulence leading to host extinction under host-parasite coevolution. BMC Evol. Biol., 15(1), pp.1–10.

Ramasamy, R., 2014. Zoonotic malaria–global overview and research and policy needs. Public Health Front., 2, p.123.

Rodrigues T, Lopes F, Moreira R. Inhibitors of the mitochondrial electron transport chain and de novo pyrimidine biosynthesis as antimalarials: The present status. Curr Med Chem. 2010;17(10):929–956.

Roth, E.J., Calvin, M.C., Max-Audit, I., Rosa, J. and Rosa, R., 1988. The enzymes of the glycolytic pathway in erythrocytes infected with Plasmodium falciparum malaria parasites. Blood (1988) 72 (6): 1922–1925

Schuster, F.L., 2002. Cultivation of Plasmodium spp. Clin. Microbiol. Rev., 15(3), pp.355–364.

Srivastava, I.K., Rottenberg, H. and Vaidya, A.B., 1997. Atovaquone, a broad spectrum antiparasitic drug, collapses mitochondrial membrane potential in a malarial parasite. J. Biol. Chem., 272(7), pp.3961–3966.

Vaidya, A.B. and Mather, M.W., 2009. Mitochondrial evolution and functions in malaria parasites. Annu. Rev. Microbiol., 63, pp.249–267.

White, N.J., 2008. The role of anti-malarial drugs in eliminating malaria. Malar. J., 7(1), pp.1–6.

Wilairatana, P., Masangkay, F.R., Kotepui, K.U., Milanez, G.D.J. and Kotepui, M., 2021. Prevalence and characteristics of malaria among COVID-19 individuals: A systematic review, meta-analysis, and analysis of case reports. Plos Negl. Trop. Dis., 15(10), p.e0009766.

